# Phylogenomic analysis of *Clostridiodes difficile* ribotype 106 strains reveals novel genetic islands and emergent phenotypes

**DOI:** 10.1101/2020.02.27.968479

**Authors:** Bryan Angelo P. Roxas, Jennifer Lising Roxas, Rachel Claus-Walker, Anusha Harishankar, Asad Mansoor, Farhan Anwar, Shobitha Jillella, Alison Williams, Jason Lindsey, Sean P. Elliott, Kareem W. Shehab, V.K. Viswanathan, Gayatri Vedantam

## Abstract

**Background:** *Clostridioides difficile* RT106 has emerged as a dominant molecular type in the USA in recent years, but the underlying factors contributing to its predominance remain undefined. As part of our ongoing *C. difficile* infection (CDI) surveillance in Arizona, we monitored RT106 frequency and characterized the genomic and phenotypic properties of the recovered isolates.

**Results:** From 2015-2018, RT106 was the second-most prevalent molecular type isolated from CDI patients in our surveillance. A representative RT106 strain displayed robust virulence and 100% lethality in the hamster model of acute CDI. We identified a unique 46 KB genomic island (GI1) in all RT106 strains, including those in public databases. GI1 was not found in its entirety in any other *C. difficile* clade, or indeed in any other microbial genome; however, smaller segments were detected in select *Enterococcus faecium* strains. Molecular clock analyses suggest that GI1 was horizontally acquired and sequentially assembled over time. Consistent with the presence of genes encoding homologs of VanZ and a SrtB-anchored collagen-binding adhesin in GI1, all tested RT106 strains had increased teicoplanin resistance and a majority displayed collagen-dependent biofilm formation. Two additional genomic islands (GI2 and GI3) were also present in a subset of RT106 strains. All three islands have features of mobile genetic elements and encode several putative virulence factors.

**Conclusions:** Consistent with the known genetic plasticity of *C. difficile*, strains belonging to the RT106 clade harbor unique genetic islands. Correspondingly, emergent phenotypic properties may contribute to the relatively rapid shifts of strain distribution in patient populations.

## BACKGROUND

The Gram-positive and spore-forming anaerobic bacterium *Clostridioides difficile* (formerly named *Clostridium difficile*) is a leading cause of antibiotic-associated diarrhea that may be self-limiting, or progress to severe and fulminant (pseudomembranous) colitis or toxic megacolon (1-4). There has been an increased incidence of *C. difficile* infection (CDI) over the past two decades (5-8) and, in the USA, this coincides with the emergence and spread of ribotype 027 strains [also called RT027 or BI or NAP1 based on the phylogenetic test (9, 10)]. While RT027 remains the most prevalent healthcare-associated *C. difficile* ribotype, its frequency has been steadily declining (11). Multiple surveillance studies indicate a changing trend in the *C. difficile* ribotype frequency distribution, particularly the emergence of RT106 (also called Group “DH” or “NAP11”) in regions where it was previously rarely found. In 2008, RT106 was second to RT027 as the most dominant ribotype in England, and was also identified in neighboring European countries including Spain and Ireland (12-14). However, during the same period, RT106 was rarely identified elsewhere in Europe, or in the USA and Canada (15), where RT027 and RT014/020 were predominant (13, 15). By 2012, RT106 emerged as the second most dominant *C. difficile* molecular type in the ten US states participating in the Centers for Disease Control and Prevention (CDC) Emerging Infections Program (EIP) surveillance (16). From 2014-2017, RT106 replaced RT027 as the most prevalent ribotype recovered from community-associated CDIs (16-21).

Currently, Arizona is not a participant in the CDC EIP program, and no molecular typing data or epidemiological trends are available for this state. As part of an ongoing surveillance to rectify this gap in knowledge, we determined the ribotype frequency of *C. difficile* isolates recovered from patients at a tertiary University Medical Center in Tucson, Arizona between August 2015 and July 2018. Consistent with broader trends in the country, we noted increased prevalence of RT106 strains in our patient population. Since little is known about these strains (22), we focused on genomic and phenotypic characterization of all recovered RT106 isolates with the goal of identifying genetic factors contributing to the increased prevalence of this molecular type.

## RESULTS

### C. difficile RT106 is the second-most prevalent healthcare-associated molecular type in Tucson, Arizona, and RT106 isolates are virulent in an animal model of infection

From August 2015 - July 2018, we recovered 788 *C. difficile* isolates from adult patients confirmed to be CDI-positive via a PCR test (employed until February 2017) or a “two-step” GDH/EIA test [Glutamate Dehydrogenase (assesses live *C. difficile*); Enzyme Immunoassay (detects *C. difficile* glycosyltransferase toxins TcdA and TcdB)] employed from March 2017. To ensure test-result consistency, we first verified the presence of *tcdB*, the same gene assayed in the PCR test, in all samples collected from March 2017 to July 2018. Overall, 519/788 isolates contained *tcdB* or expressed EIA-detectable levels of TcdA/B. Ribotype analysis revealed a diversity of strains in the patient population, with RT027 being the most frequently isolated strain (n=144) (**Figure 1**). RT106 (n=38) was the second most frequently identified ribotype over the three-year period.

**Figure 1.**
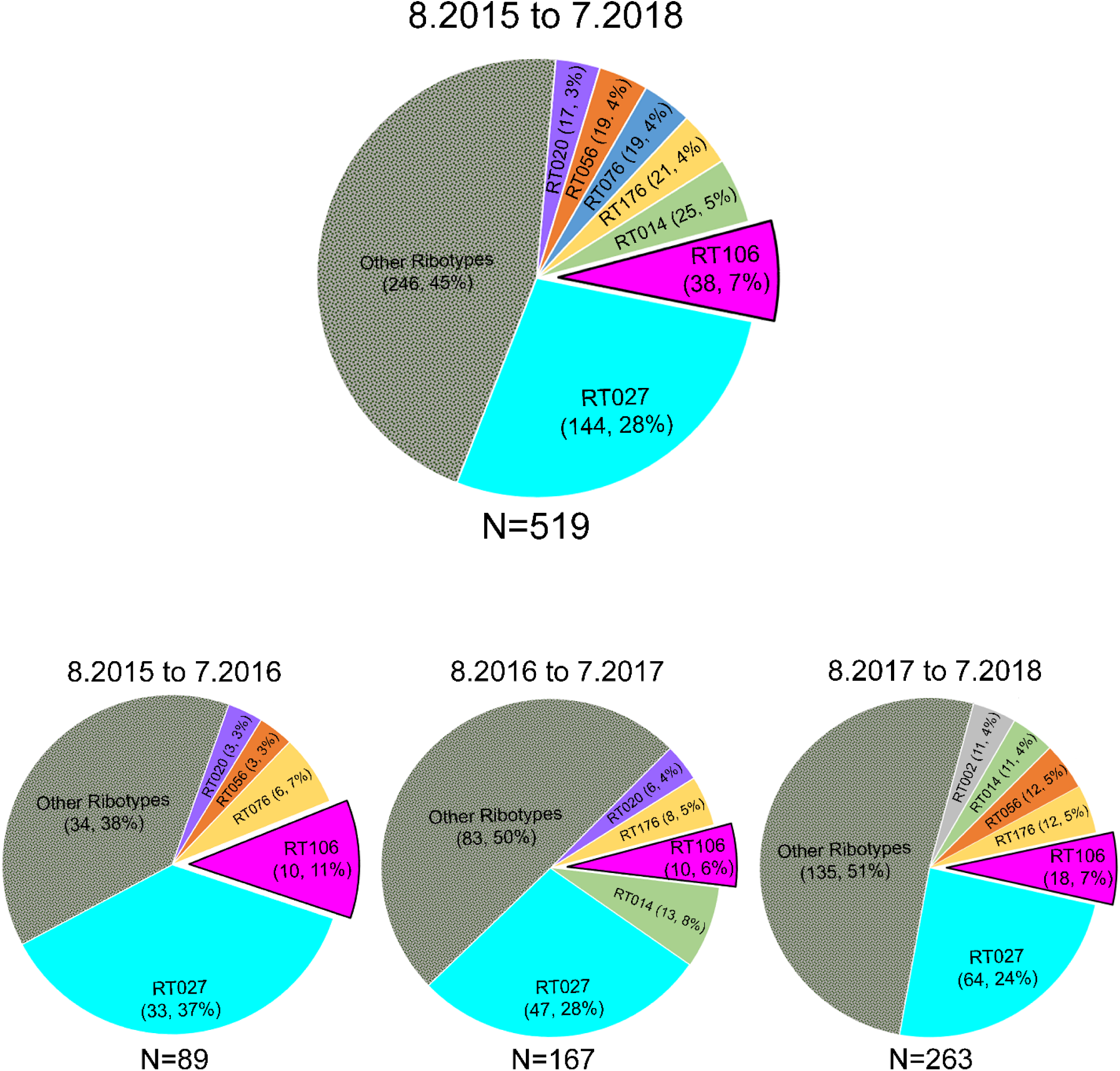
RT106 is the second most prevalent molecular type in a Tucson-area hospital. Top chart depicts ribotype distribution of 519 *tcdB* PCR-positive and/or TcdA/B ELISA-positive *C. difficile* strains from patient stool samples collected from August 2015 to July 2018 (8.2015 to 7.2018). Ribotype frequency and percent of total sample size are shown in parenthesis. Overall, RT106 is the second most frequently isolated molecular type, while RT027 is the most prevalent ribotype. Bottom charts depict ribotype distribution in 12-month periods. RT106 ranked second to RT027 as the most frequently isolated molecular type during 8.2015 to 7.2016 and 8.2017 to 7.2018. RT106 was the third most dominant ribotype during 8.2016 to 7.2017.

Prior to detailed characterization of RT106 isolates, we verified the virulence of the representative strain GV599 in the Golden Syrian hamster model of acute *C. difficile* infection. All infected animals succumbed to disease within 2-6 days of spore inoculation (**Supplemental Figure S1A**). Microscopy-based visualization of colonic tissue sections revealed classic *C. difficile* infection pathology including gross hemorrhage, epithelial erosion and inflammatory infiltrates (**Supplemental Figure S1B**).

### RT106 strains group monophyletically, and harbor one clade-specific, and up to two other, novel genetic elements

Whole genome sequencing was performed on all 38 RT106 strains recovered in our surveillance, and data were compared to 1425 publicly available *C. difficile* strain sequences. Based on single nucleotide polymorphism (SNP) analyses (23, 24), the strains were not clonal, and the two closest-related isolates (GV597 and GV753) were divergent by 113 SNPs. Overall, RT106 genomes were most-closely related to RT002 strains (25).

Our 38 RT106 strains mapped closely to 33 previously sequenced RT106 strains from pediatric patients (26, 27) (**Figure 2B and 2C**). Twenty-three other *C. difficile* strains of unknown ribotype also claded strongly with RT106. We performed *in silico* ribotyping on these 23 strains, and 13/23 (those with currently available closed genome sequence) generated a clear RT106 PCR fragment pattern. For an additional assessment of genome relatedness, we performed *in silico* Multi-Locus Sequence Typing (MLST) on all 94 strains; this method differentiates organisms into Sequence Types [STs; (28)]. 92/94 strains were sequence type ST42, whereas 2/94 belonged to the closely-related sequence type ST28 (29). Taken together, all 94 strains interrogated in these analyses grouped together in a distinct monophyletic RT106 clade [(**Figure 2B and 2C**);(30)].

**Figure 2.**
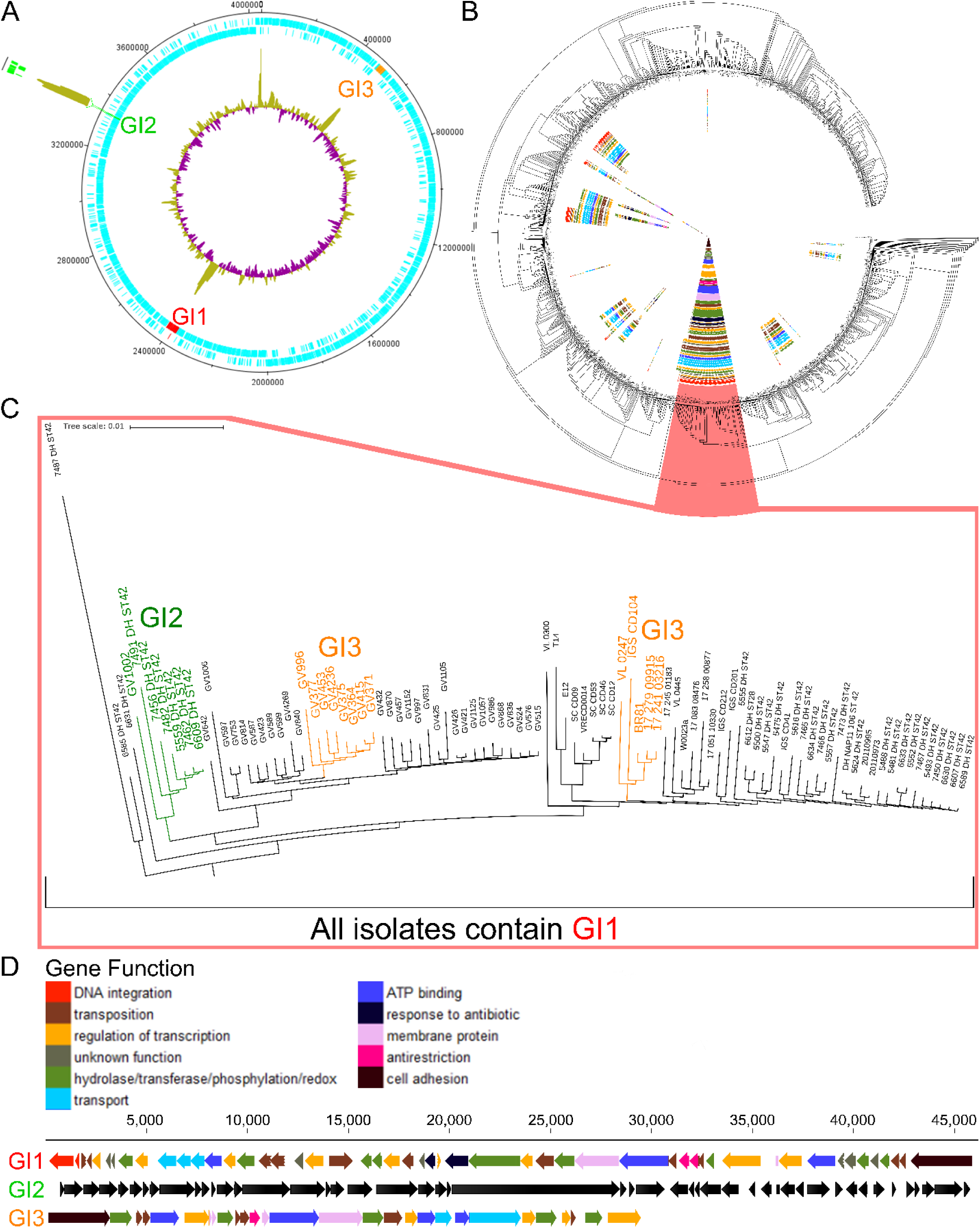
The monophyletic RT106 isolates may harbor up to three novel genomic islands. **A**, Genetic islands GI1, GI2, and GI3 associated with RT106 are at three different locations in the genome. GV364, used as a representative genome, contains GI1 and GI3. GI-2 was previously identified in pediatric RT106 strains from Chicago, Illinois (46). Insertion site (green) of genomic island 2 is shown relative to GI-1 and GI3 locations. **B**, A phylogenetic tree of 1425 publicly available *C. difficile* and 38 RT106 genome sequences shows that RT106 strains clade together in a monophyletic branch. Gene segments of GI1 were found in several strains outside the RT106 clade, but not as a complete island. **C**, Currently, RT106 clade (red box) consists of 94 strains. All 94 strains within the RT106 clade harbor the complete GI1. GI-2 is present in 7 RT106 strains (green) (6 pediatric isolates RT106 strains from Chicago, Illinois, and GV1002 from Tucson, Arizona). Thirteen strains harboring GI3 (yellow) belong to 2 different subclades. **D**, Schematic shows gene arrangement, size and functions of GI1, GI2 and GI3.

Up to three unique genomic islands GI1, GI2 and GI3 are associated with the RT106 clade (**Figure 2A**), and GI1, a novel 46 kb element reported for the first time herein, is invariantly carried by all RT106 strains. GI2 was previously identified in RT106 strains recovered from pediatric patients (27), and its overall prevalence in the RT106 clade is 7.4% (7/94 strains). GI3 (a 29.4 kb element) prevalence is 13.8% (13/94 strains). GI1, GI2 and GI3 all have features of mobile genetic elements and contain DNA integration and transposition genes (**Supplemental Table S1** and **Supplemental Table S2**); genes predicted to encode anti-restriction modification, antibiotic-resistance and cell adhesion functions are also present in GI1 and GI3 (**Figure 2D; Supplemental Tables S1** and **S2**). All three islands display higher percentage G+C content (38%, 45% and 37% for GI1, GI2 and GI3, respectively) than the rest of the *C. difficile* genome (28-29%).

Currently, the 46 kb GI1 appears to be uniquely and specifically associated with RT106 (**Figure 2B &2C**), and all sequenced strains belonging to this clade (38 from this study and 56 others identified in publicly available databases) harbor a complete GI1 island. GI1 has 99.91% pairwise identity among strains (100% GI1 identity in 48 strains; 44 strains with 1-2 SNPs; 2 strains with >3 SNPs). Fragments of GI1 were, however, detected in some non-RT106 strains. GI2, previously identified in pediatric RT106 isolates (27), is present in only 1/38 adult RT106 strains from our surveillance (**Figure 2C**); we also identified this island in the non-RT106 strain Y358 (GCF_00451525.2). The 29.4 kb GI3 is present in 8/38 of our adult RT106 strains, as well as 5 other RT106 isolates in publicly available databases (**Figure 2C**). We also identified GI3 in one non-RT106 strain (VRECD0053, GCF_900164815.1).

### The 46 kb genomic island 1 is unique to RT106/ST42/ST28 strains

BLAST analysis of the 46 kb GI1 against 1425 publicly available *C. difficile* genome sequences at the NCBI database resulted in the identification of 265 *C. difficile* strains that contain either segments of or the entire genomic island (>7.7 kb, 98% identity). We concomitantly performed in silico MLST analysis to determine the respective sequence types, and then generated a phylogenetic tree using CVtree Version 3.0 (30). GI1-related genes found in each strain were annotated based on gene function. Only RT106/ST42/ST28 strains harbor the complete 46 kb GI1, while other ST strains included in this analysis contain only shorter segments of the genomic island (**Figure 3**).

**Figure 3.**
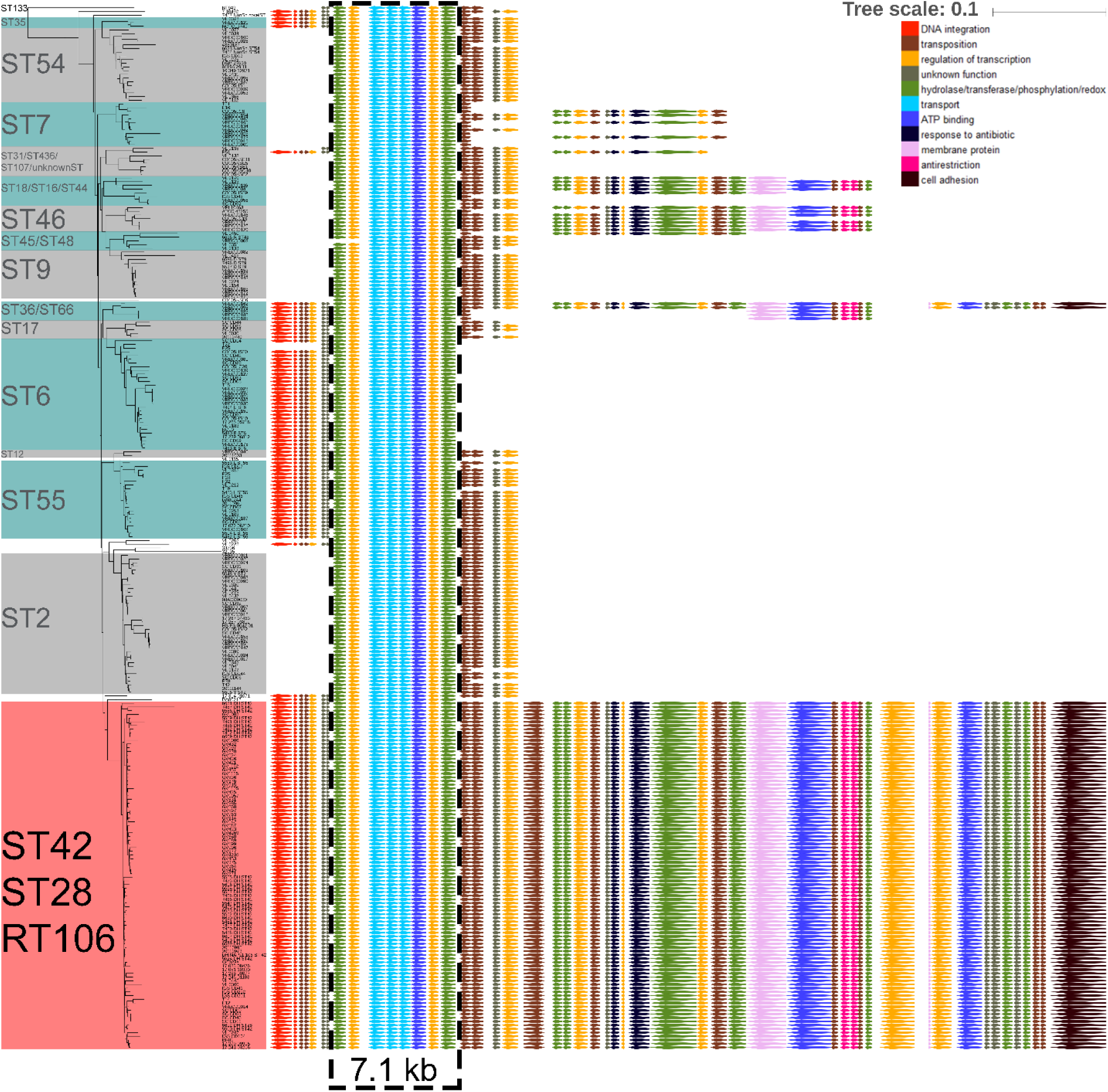
RT106 strains harbor a complete and unique 46 kb genomic island 1. A phylogenetic tree of *C. difficile* strains that carry segments (>7.7kb, 98% identity) of the GI1 identified in RT106 was generated. GI1 is drawn to scale on the right to illustrate regions present in different sequence types (ST). The complete 46 kb GI1 is present in RT106/ST28/ST42. Genes were colored based on functional categories from gene ontology (GO) analysis. A 7.1 kb region carried by all the strains (grey dashed box) was used for determining progenitor STs of the element in the molecular clock analysis in Figure 4.

A 7.1 kb gene segment (demarcated within a hatched black box; **Figure 3**) is common to all MLST sequence type strains shown. SNP analysis was performed on the 7.1 kb gene segment to generate a molecular clock of GI1 via Mega-X (31) using maximum likelihood (ML) approach (**Figure 4**). The molecular clock revealed gradual and progressive acquisition of gene elements in different strains, finally leading to an intact GI1. ST133, which branches most distantly from RT106, contained the least number of GI1-associated genes as opposed to STs branching closer to RT106/ST42/28.

**Figure 4.**
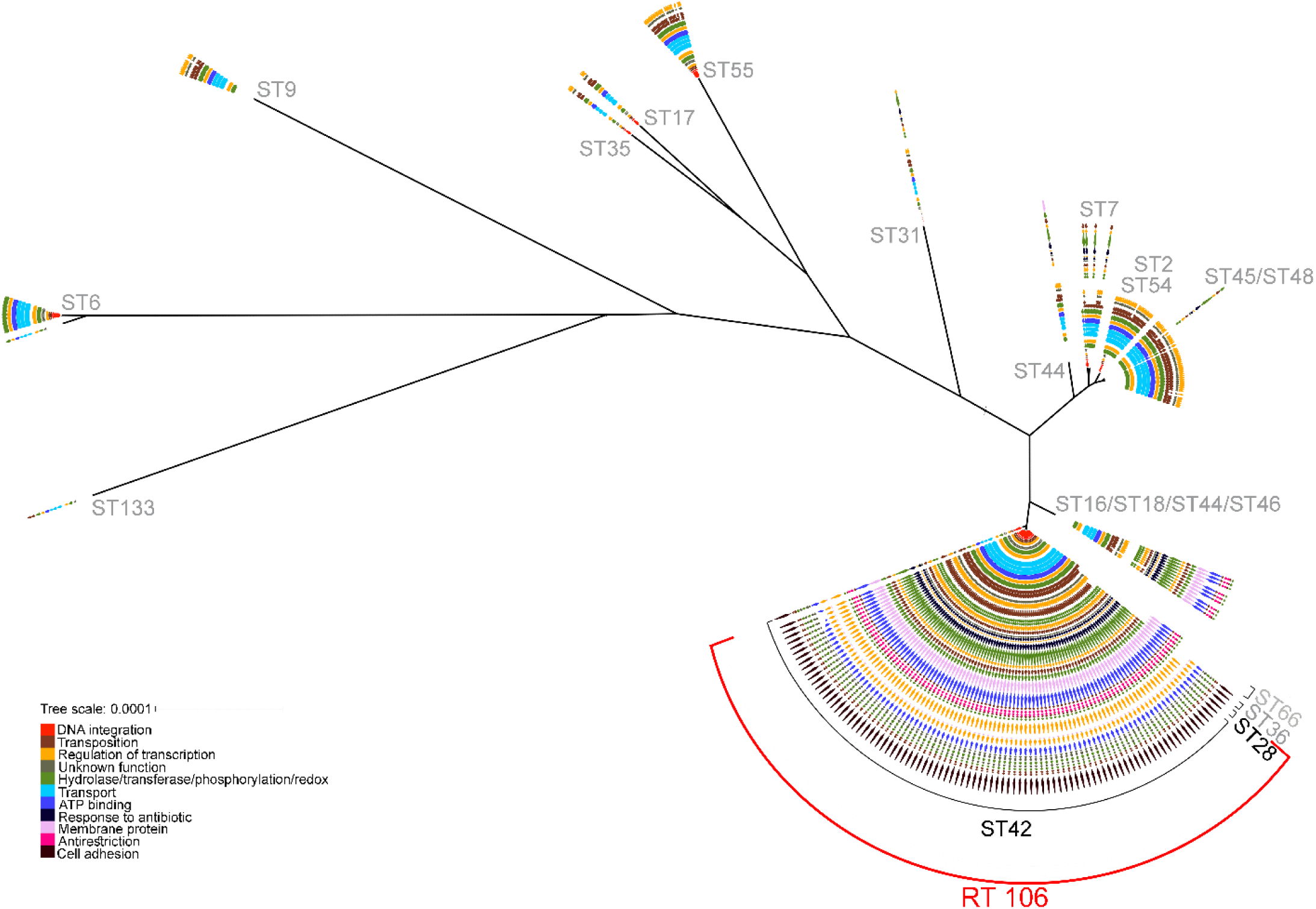
Molecular clock analysis reveals organization of the 46 kb genomic island 1 via acquisition of distinct sub-elements. Maximum likelihood (ML) molecular clock analysis of *C. difficile* strains that carry the 46 kb GI1. Evolutionary genetic analysis was performed on the 7.1 kb region highlighted in grey dashed box in Figure 3. Tree generated using ML and branch lengths were adjusted using Tajima’s relative rate test (molecular clock).

To definitively verify that the various GI1 segments were acquired via horizontal transfer, we used the seven housekeeping genes utilized in MLST characterization to establish genetic relatedness of strains harboring the core 7.1 kb GI1 fragment (**Supplemental Figure S2**). ST28, a sequence type that is included within the RT106 clade, is closely related to ST16, ST18, ST44 and ST46 based on sequence similarity of the both core GI1 island 1 fragment (**Figure 4)** and the seven MLST gene loci (**Supplemental Figure S2**). However, this is not the case with the more predominant sequence type, ST42, within the RT106 clade. ST42 is closer to ST48 and ST7 based on the seven housekeeping genes (**Supplemental Figure S2**), and yet these ST strains map distantly in the core GI1-based molecular clock (**Figure 4**).

GI1 is not found in any other bacteria. However, two regions (8.4 kb and 13.7 kb) within GI1 were detected in other enteric bacteria (**Figure 5**). The 13.7 kb gene segment was found in *Enterococcus faecium* EnGen0312 UAA407 at 99% sequence identity, while the 8.4 kb gene segment occurs in the same gene order but with some sequence plasticity in *Enterococcus faecium* EnGen0312 UAA407, *Anaerostipes hadrus* BPB5-Raf3-2-5, *Clostridioides sporogenes* YH-Raf3-2-5 and *Roseburia intestinalis* M50/1 strains (89.8%, 90.4%, 90.5% and 92.2% DNA sequence identity, respectively).

**Figure 5.**
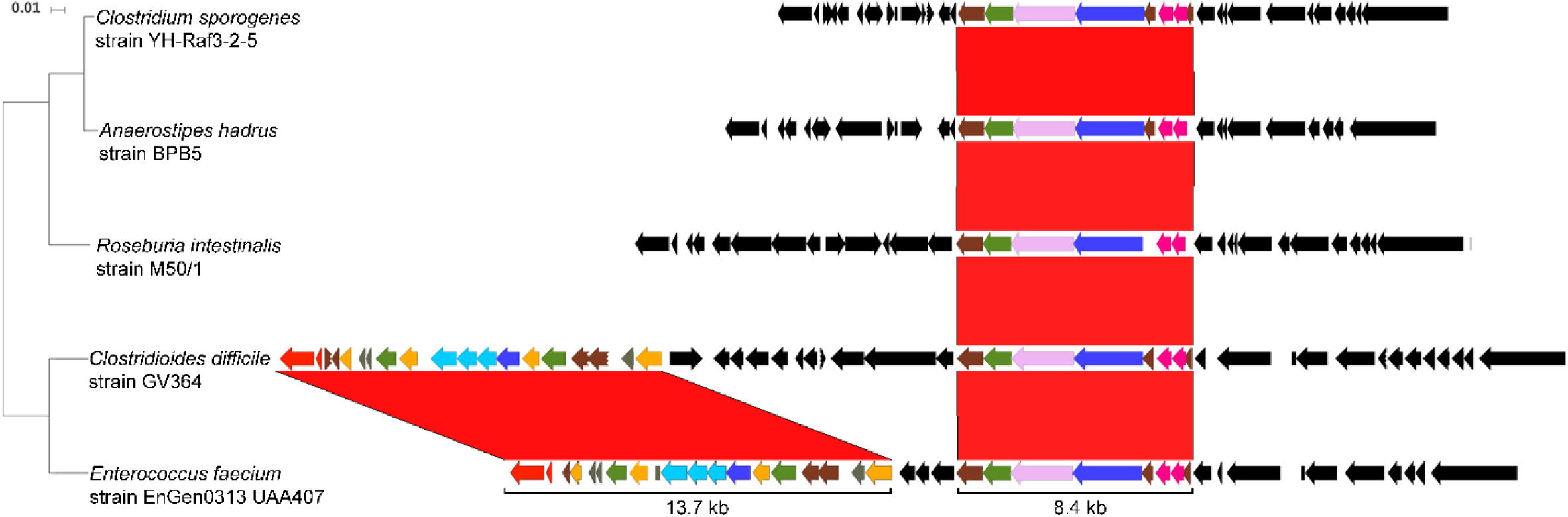
Human commensal microbiota may contribute to the acquisition of the 46 kb genomic island 1. The complete 46 kb GI1 is not present in any other microbial genome nor plasmid sequence, but two gene segments (8.4 kb and 13.7 kb) within the island are found in other human enteric bacteria. The 8.4 kb gene segment is present in *Enterococcus faecium* EnGen0312 UAA407, *Anaerostipes hadrus* BPB5-Raf3-2-5, *Clostridioides sporogenes* YH-Raf3-2-5 and *Roseburia intestinalis* M50/1 strains (89.8%, 90.4%, 90.5% and 92.2% DNA sequence identity, respectively). *E. faecium* also harbors a 13.7 kb gene segment at 99% sequence identity. These two gene segments are found in *E. faecium* as part of a 36 kb genomic element. RT106 strains do not carry this 36 kb genomic element, but other *C. difficile* strains (VL0228 and 17-314-01071) strains have the identical 36 kb genomic element. Comparative genomic analyses were performed using BLASTN and results were viewed using Artemis Release 17.0.1.

### Phenotypic Characterization of RT106 isolates

Clade-specific properties, including those conferred by genes within GI1 could explain the emergence and spread of RT106 strains. We, therefore, assessed various virulence-associated phenotypes including antibiotic susceptibility, motility, toxin production, biofilm production and adhesion to collagen on the first 21 of the 38 RT106 strains chronologically obtained from our clinical surveillance.

### RT106 strains display variable antibiotic susceptibility, with some isolates displaying multi-drug resistance

We determined the susceptibility of RT106 isolates to the antibiotics cefotaxime, vancomycin, erythromycin, clindamycin, levofloxacin, moxifloxacin, metronidazole, and tetracycline. All isolates were resistant to cefotaxime (minimum inhibitory concentration (MIC) >32 mcg/ml), but susceptible to vancomycin, metronidazole, and tetracycline (**Table 2**). 18/21 strains had intermediate resistance to clindamycin (MIC = 4-6 mcg/ml). Three isolates (GV371, GV423, GV432) were highly resistant to erythromycin (MIC >256 mcg/ml). Clindamycin and erythromycin belong to the macrolide-lincosamide-streptogramin B (MLS_B_) group of protein synthesis inhibitors. MLS_B_ resistance in *C. difficile* has been associated with the acquisition of *erm* genes (32) or nucleotide substitution (C→T) at position 656 within the 23S rDNA (33). None of the RT106 strains harbor the *erm* genes, while only GV415 had the 23S rDNA 656C>T substitution; however, GV415 has low-level resistance to clindamycin (MIC = 4 mcg/ml).

**Table 1.** List of RT106 Strains

| Strain | Isolation Year | Clinical Test<br>(PCR or GDH/EIA Result) | Genome Size<br>(bases) | GC Content (%) | Phenotypic Characterization<br>Performed? (Yes or No) |
| --- | --- | --- | --- | --- | --- |
| GV364 | 2015 | PCR: + | 4043408 | 28.4 | Yes |
| GV371 | 2015 | PCR: + | 4062530 | 28.4 | Yes |
| GV375 | 2016 | PCR: + | 4045723 | 28.4 | Yes |
| GV377 | 2016 | PCR: + | 4081010 | 28.5 | Yes |
| GV415 | 2016 | PCR: + | 4052544 | 28.4 | Yes |
| GV421 | 2016 | PCR: + | 4012174 | 28.3 | Yes |
| GV423 | 2016 | PCR: + | 4050785 | 28.4 | Yes |
| GV425 | 2016 | PCR: + | 4014240 | 28.3 | Yes |
| GV426 | 2016 | PCR: + | 4013299 | 28.3 | Yes |
| GV432 | 2016 | PCR: + | 4051197 | 28.5 | Yes |
| GV453 | 2016 | PCR: + | 4126224 | 28.5 | Yes |
| GV457 | 2016 | PCR: + | 4067987 | 28.4 | Yes |
| GV515 | 2017 | GDH/EIA: +/+ | 4030448 | 28.3 | Yes |
| GV524 | 2017 | GDH/EIA: -/- | 4022675 | 28.3 | Yes |
| GV576 | 2017 | GDH/EIA: +/- | 4027239 | 28.3 | Yes |
| GV587 | 2017 | GDH/EIA: +/- | 4051442 | 28.4 | Yes |
| GV589 | 2017 | GDH/EIA: +/+ | 4061628 | 28.4 | Yes |
| GV597 | 2017 | GDH/EIA: +/- | 4059478 | 28.4 | Yes |
| GV599 | 2017 | GDH/EIA: +/+ | 4083315 | 28.5 | Yes |
| GV642 | 2017 | GDH/EIA: +/- | 4103497 | 28.4 | Yes |
| GV753 | 2017 | GDH/EIA: +/+ | 4094464 | 28.4 | Yes |
| GV814 | 2017 | GDH/EIA: +/+ | 4066154 | 28.5 | No |
| GV831 | 2017 | GDH/EIA: +/+ | 4048397 | 28.5 | No |
| GV836 | 2017 | GDH/EIA: +/+ | 4039872 | 28.4 | No |
| GV840 | 2017 | GDH/EIA: +/+ | 4066224 | 28.4 | No |
| GV868 | 2018 | GDH/EIA: +/+ | 4026007 | 28.4 | No |
| GV870 | 2018 | GDH/EIA: +/+ | 4080981 | 28.8 | No |
| GV962 | 2018 | GDH/EIA: +/+ | 4086548 | 28.4 | No |
| GV973 | 2018 | GDH/EIA: +/+ | 4112429 | 28.6 | No |
| GV986 | 2018 | GDH/EIA: +/+ | 4037148 | 28.4 | No |
| GV996 | 2018 | GDH/EIA: +/+ | 4112462 | 28.7 | No |
| GV997 | 2018 | GDH/EIA: +/+ | 4053485 | 28.5 | No |
| GV1002 | 2018 | GDH/EIA: +/+ | 4097153 | 29 | No |
| GV1006 | 2018 | GDH/EIA: +/+ | 4412809 | 29.2 | No |
| GV1057 | 2018 | GDH/EIA: +/+ | 4038504 | 28.5 | No |
| GV1105 | 2018 | GDH/EIA: +/+ | 4070520 | 28.7 | No |
| GV1125 | 2018 | GDH/EIA: +/+ | 4028672 | 28.4 | No |
| GV1152 | 2018 | GDH/EIA: +/+ | 4055778 | 28.4 | No |

**Table 2.** Antibiotic susceptibility profiles of RT106 clinical isolates (this study)

| Strain | Minimum Inhibitory Concentration (MIC) |  |  |  |  |  |  |  |  |
| --- | --- | --- | --- | --- | --- | --- | --- | --- | --- |
|  | Teicoplanin | Cefotaxime | Clindamycin | Erythromycin | Levofloxacin | Moxifloxacin | Tetracycline | Vancomycin | Metronidazole |
|  | 0.016-256 mcg/mL | 0.002-32 mcg/mL | 0.016-256 mcg/mL | 0.016-256 mcg/mL | 0.002-32 mcg/mL | 0.002-32 mcg/mL | 0.016-256 mcg/mL | 0.016-256 mcg/mL | 0.016-256 mcg/mL |
| GV371 | 0.125 | > 32 ⚡ | 4 ⚡ | >256 □ | 4 | 3 | 0.50 | 0.75 | 0.50 |
| GV423 | 0.125 | > 32 ⚡ | 6 ⚡ | >256 □ | 4 | 3 | 0.50 | 1.00 | 0.50 |
| GV432 | 0.125 | > 32 ⚡ | 4 ⚡ | >256 □ | 4 | 2 | 0.09 | 1.00 | 0.38 |
| GV597 | 0.094 | > 32 ⚡ | 4 ⚡ | 2 | 12 □ | 4 ⚡ | 0.38 | 0.75 | 0.38 |
| GV453 | 0.125 | > 32 ⚡ | 4 ⚡ | 1.50 | 4 | 4 ⚡ | 0.50 | 1.00 | 0.50 |
| GV587 | 0.125 | > 32 ⚡ | 6 ⚡ | 1 | 4 | 4 ⚡ | 0.50 | 0.75 | 0.38 |
| GV642 | 0.125 | > 32 ⚡ | 4 ⚡ | 1.50 | 4 | 6 ⚡ | 0.38 | 0.75 | 0.50 |
| GV364 | 0.125 | > 32 ⚡ | 4 ⚡ | 1 | 6 | 2 | 0.50 | 1.00 | 0.50 |
| GV375 | 0.125 | > 32 ⚡ | 4 ⚡ | 1 | 6 | 2 | 0.38 | 0.50 | 0.75 |
| GV377 | 0.125 | > 32 ⚡ | 4 ⚡ | 2 | 4 | 3 | 0.38 | 0.75 | 0.75 |
| GV415 | 0.125 | > 32 ⚡ | 4 ⚡ | 1 | 6 | 3 | 0.50 | 1.00 | 0.50 |
| GV421 | 0.125 | > 32 ⚡ | 4 ⚡ | 1 | 4 | 2 | 0.38 | 0.75 | 0.75 |
| GV425 | 0.125 | > 32 ⚡ | 6 ⚡ | 0.75 | 4 | 3 | 0.06 | 0.75 | 0.19 |
| GV426 | 0.094 | > 32 ⚡ | 4 ⚡ | 1.50 | 6 | 3 | 0.50 | 0.75 | 0.50 |
| GV524 | 0.125 | > 32 ⚡ | 4 ⚡ | 0.75 | 4 | 3 | 0.50 | 0.75 | 0.38 |
| GV576 | 0.125 | > 32 ⚡ | 4 ⚡ | 1 | 4 | 3 | 0.38 | 0.75 | 0.50 |
| GV589 | 0.094 | > 32 ⚡ | 4 ⚡ | 1 | 4 | 2 | 0.38 | 0.75 | 0.50 |
| GV753 | 0.125 | > 32 ⚡ | 6 ⚡ | 1 | 4 | 3 | 0.38 | 0.75 | 0.50 |
| GV599 | 0.094 | > 32 ⚡ | 3 | 1 | 4 | 3 | 0.38 | 0.50 | 0.38 |
| GV457 | 0.094 | > 32 ⚡ | 3 | 1 | 4 | 3 | 0.13 | 0.75 | 0.38 |
| GV515 | 0.125 | > 32 ⚡ | 3 | 1 | 4 | 2 | 0.38 | 1.00 | 0.50 |
| 630 | 0.064 | > 32 ⚡ | >256 □ | >256 □ | 4 | 3 | 64 □ | 1.50 | 0.25 |
| VPI | 0.032 | > 32 ⚡ | 4 ⚡ | 1 | 3 | 1 | 0.38 | 1.00 | 0.25 |
| T-7 | 0.064 | > 32 ⚡ | 3 | 1 | 4 | 3 | 0.50 | 1.00 | 0.50 |
| BI-1 | 0.064 | > 32 ⚡ | 2 | 1 | 4 | 2 | 0.38 | 1.50 | 0.25 |
| CLSI Breakpoints<br>(mcg/mL) | NA | ≥ 64 (Resistant) | ≥ 8 (Resistant) | NA | NA | ≥ 8 (Resistant) | ≥ 16 (Resistant) | NA | ≥ 32 (Resistant) |
|  | NA | 32 (Intermediate Resistance) | 4 (Intermediate Resistance) | NA | NA | 4 (Intermediate Resistance) | 8 (Intermediate Resistance) | NA | 16 (Intermediate Resistance) |
|  | NA | ≤ 16 (Susceptible) | ≤ 2 (Susceptible) | NA | NA | ≤ 2 (Susceptible) | ≤ 4 (Susceptible) | NA | ≤ 8 (Susceptible) |
| EUCAST Breakpoints<br>(mcg/mL) | NA | NA | NA | NA | NA | NA | NA | >2 (Resistant) | >2 (Resistant) |
|  | NA | NA |  | NA | NA | NA | NA | ≤ 2(Susceptible) | ≤ 2(Susceptible) |

Notes:
- ⚡ denotes that strain is moderately resistant to specific antibiotics - denotes that strain is highly resistant to specific antibiotics

All RT106 isolates, except GV597, were susceptible to the fluoroquinolone levofloxacin. GV597, GV453, GV587, and GV642 had intermediate resistance to the fluoroquinolone moxifloxacin (MIC = 4-6 mcg/ml). However, these fluoroquinolone-resistant RT106 isolates do not encode mutations in GyrA (Thr-82-Ile, Thr-82-Val, Asp-71-Val, Asp-81-Asn and Ala-118-Thr) or GyrB (Asp-426-Val, Asp-426-Asn, Arg-447- Leu, Arg-447-Lys, Ser-366-Ala and Ser-416-Ala) associated with fluoroquinolone resistance (34-38).

GI1 harbors a gene encoding a VanZ family protein (locus ID FE556_12215; **Supplemental Table S1**) previously implicated in teicoplanin resistance. Consistent with this, all RT106 isolates exhibit modest increase in resistance to teicoplanin compared to reference strains (T7, BI-1, 630, VPI; **Table 2**); the teicoplanin CLSI and EUCAST breakpoint values for *C. difficile* have not been established. Cultivation of RT106 strains in sub-inhibitory concentration (MIC) of teicoplanin (0.0125 mcg/mL) resulted in increased teicoplanin resistance in 7/21 strains (**Supplemental Table S3**).

### RT106 strains display collagen-dependent biofilm formation

Biofilm formation could facilitate intestinal colonization and persistence, and possibly contribute to recurrence. RT106 strains display variable biofilm densities on abiotic plastic surface (**Figure 6A**). Since GI1 encodes a putative SrtB-anchored collagen-binding adhesin (locus ID FE556_11350; **Supplemental Table S1**), we tested the ability of RT106 strains to form biofilms on type I and type III collagen, the major collagen types present in the extracellular matrix of normal human intestines (39).

**Figure 6.**
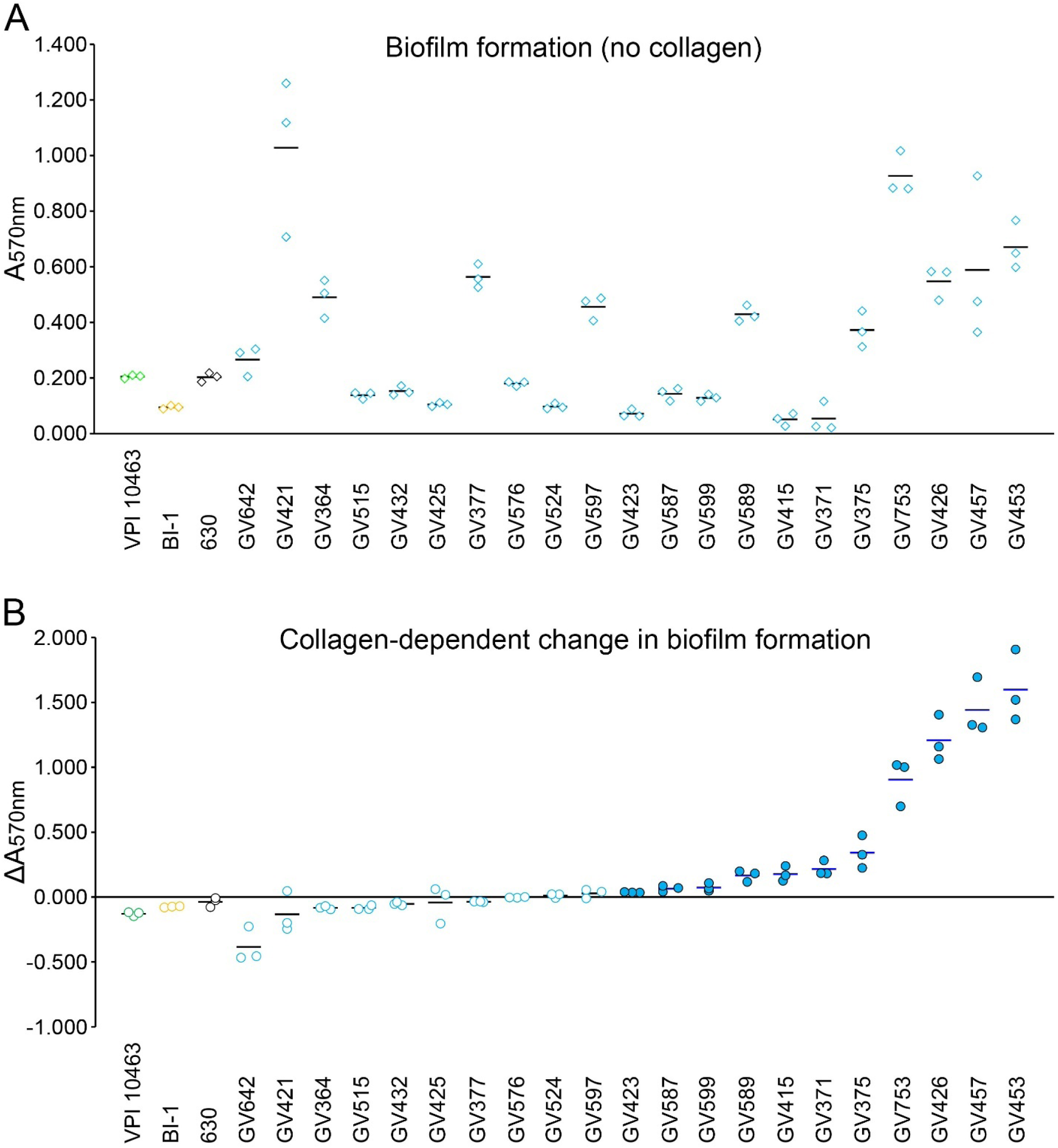
Clinical RT106 isolates display collagen-dependent biofilm formation. **A**, 21 clinical RT106 strains (blue circles) and 3 non-RT106 toxigenic *C. difficile* strains (VPI, BI-1, and 630 designated as green, yellow and black circles, respectively) were cultured in uncoated or collagen-coated (combined types I and III) plastic wells for 72 hours. Biofilm assay was performed by measuring absorbance at λ=570 nm (A_570nm_) of solubilized crystal violet-stained biofilms. RT106 strains displayed variable levels of biofilm on abiotic plastic wells. **B**, Relative changes in biofilm densities (ΔA_570nm_) were determined by comparing A_570nm_ of crystal violet-stained biofilms formed on human collagen (combined types I and III) vs. on uncoated plastic wells. Student’s t test was performed to compare mean absorbance at 570nm of biofilm formed by each strain on collagen-coated vs. uncoated wells. Filled blue circles denote P_value_ < 0.05. No difference in biofilm formation was observed when the reference *C. difficile* 630, BI-1 and VPI strains were cultured on wells with or without collagen. One-sample one-tailed T-test was performed to determine whether the group of RT106 strains displayed denser biofilms on collagen-coated wells (H_alt_: mean ΔA_570nm_>0; H_0_: mean ΔA_570nm_=0; P_value_=0.02038).

Biofilm densities of the non-RT106 toxigenic *C. difficile* strains BI1, 630 and VPI did not increase in the presence of collagen (**Figure 6B)**. Eleven RT106 strains displayed collagen-dependent increase in biofilm formation when cultured on wells coated with both type I and type III human collagen. Overall, RT106 strains, as a group, have increased likelihood of displaying collagen-dependent biofilm formation.

We also interrogated the ability of the strains to form biofilms on either human Type I or Type III collagen. Although some RT106 strains showed increased biofilm formation on either collagen type (6 to human type I collagen; 5 to human type III collagen), the RT106 strain group did not show collagen-dependent biofilm formation when only one collagen type was used for collagen coating. Curiously, GV453, GV457 and GV467 showed synergistic increase in biofilm formation to human types I and III collagen.

We also tested the ability of the 21 RT106 strains to form biofilms on rat type 1 collagen and found that ten strains formed denser biofilms on rat collagen compared to uncoated plastic wells (**Supplemental Figure S4**). GV425, GV426, GV432, GV453 and GV457 consistently formed denser biofilms on human and rat type I collagen compared to uncoated wells.

### RT106 strains are variably motile

Flagella-dependent motility influences virulence of many pathogens. All RT106 isolates tested, except GV375, GV415 and GV426, were motile (**Figure 7**). We analyzed the genome of the non-motile RT106 strains for mutations in flagella-associated genes. In *C. difficile* 630 strain, flagella-associated genes are found in the F1 and F3 loci (40, 41). F1 and F3 loci are highly conserved in RT106; therefore, the nonmotile phenotype observed for GV375, GV415 and GV426 may possibly result from alterations in expression and/or post-translational modifications.

**Figure 7.**
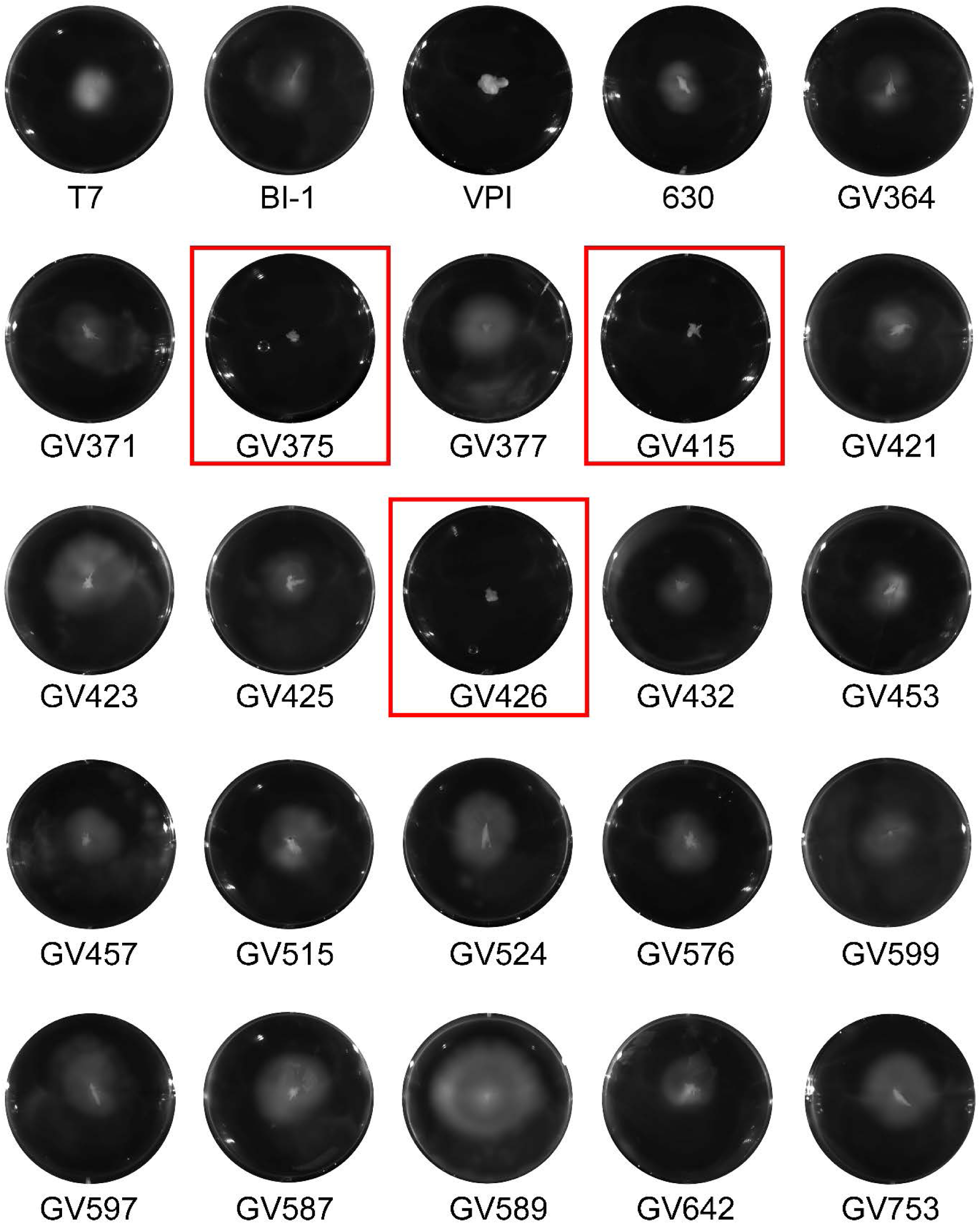
Clinical RT106 isolates are variably motile. All 21 clinical RT106 isolates, except GV375, GV415 and GV426 (red box), were motile in BHI soft agar. Motile (T7, BI-1, 630) and non-motile (VPI) reference strains are shown.

### *Most RT106 strains* are robust toxin-producers

Toxigenic *C. difficile* produce up to two related glucosylating toxins, toxin A (TcdA) and toxin B (TcdB), which are encoded on the pathogenicity locus (PaLoc) (42, 43). Genome analysis of RT106 isolates revealed that all strains harbor the complete PaLoc and the gene for the TcdB1, instead of the highly toxigenic TcdB2 variant associated with select ribotypes including RT027 (44, 45). We quantified secreted toxin, and observed that all RT106 strains, except GV457 and GV423, produced detectable TcdA/TcdB levels (**Figure 8**). Nine RT106 isolates expressed TcdA/TcdB at levels comparable to the reference strain 630, while ten RT106 strains had similar (4/10) or higher (6/10) TcdA/TcdB levels compared to the RT027 strain BI-1.

**Figure 8.**
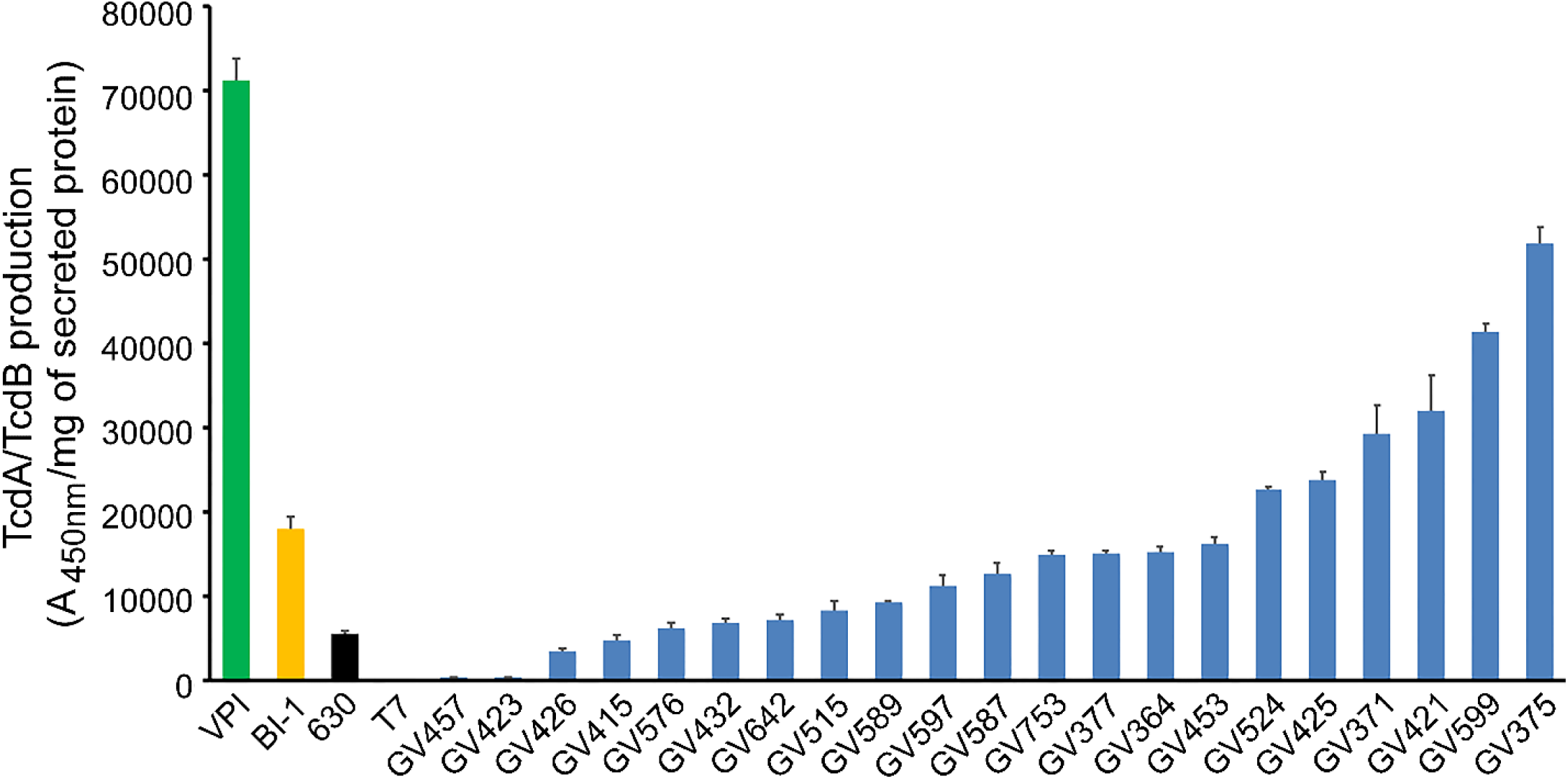
Most clinical RT106 isolates are robust toxin producers. All RT106 samples, except for GV457 and GV423, secrete similar or greater TcdA/TcdB levels compared to *C. difficile* 630 strain. Four strains (GV753, GV377, GV364, GV453) produced similar TcdA/TcdB levels as the BI-1 reference strain. Six strains (GV524, GV425, GV371, GV421, GV599, GV375) secrete more TcdA/TcdB compared to BI-1. TcdA/TcdB levels secreted after 72-hour culture in BHI broth were normalized to mg of total secreted proteins. Mean A_450nm_/mg of secreted protein and standard deviation are shown. Image is representative of two independent TcdA/TcdB ELISA assays with three sample replicates per condition.

## DISCUSSION

Consistent with broader trends in the United States, RT106 has emerged as the second leading ribotype from healthcare-associated cases in Southern Arizona (15-21). Our genotypic and phenotypic characterization of multiple RT106 strains, along with the recent studies by Kociolek et al, represents an initial foray into defining key virulence properties of this clade (27, 29, 46).

The factors contributing to the emergence and expansion of RT106 strains are presently undefined, but they appear to be distinct from those postulated for the healthcare- and US-dominant RT027 clade. First, the enhanced ability of RT027 strains to utilize trehalose, a sugar increasingly used in food products since the early 2000, may have provided a selective advantage for this clade (although this has recently been disputed) (47, 48). None of the 94 sequenced RT106 genomes harbor the Leu-1721-Ile substitution in the TreR repressor or the four-gene insertion sequence that allow RT027 and RT078 strains, respectively, to grow on low levels of trehalose (47). Still, our studies do not rule out unique sugar- or carbon source-utilization capabilities of RT106 strains.

Second, DNA gyrase mutations conferring fluoroquinolone resistance may have contributed to the emergence and spread of RT027 strains (35). While RT106 isolates from the United Kingdom were highly resistant to moxifloxacin, those from North American surveillance studies, including ours, were mostly susceptible to fluoroquinolones (Table 2) (12, 49-51). Thus, fluoroquinolone resistance does not explain their emergence and spread in the United States.

Third, the PaLoc region of RT027 strains displays several key differences relative to the historic strain 630 (RT012) (52). These include a point mutation in *tcdC* (though not in all isolates) that results in a truncated version of the anti-sigma factor TcdC, and expression of a variant of toxin B (TcdB2). TcdB2 has enhanced ability to enter host cells, is more cytotoxic, and exhibits wider tissue tropism (44, 45). In contrast to RT027 strains, the PaLoc of RT106 strains is 100% identical to 630 (53-56); thus, these strains encode full-length TcdC and express the TcdB1 toxin variant. Both RT027 and RT106 isolates produce variable amounts of TcdA/TcdB. Also, unlike 630 and RT106 strains, RT027 strains encode the binary toxin. Thus, toxin variations seem to be an unlikely driving force for the spread of RT106 strains.

Detailed genome sequence analyses, however, suggest that the acquisition of novel genetic islands may be a contributor to RT106 emergence. All sequenced RT106 strains harbor a unique 46 kb genomic island (GI1) with a distinct GC content suggestive of horizontal acquisition. GI1 possesses several gene attributes that may confer competitive advantage to the RT106 clade. It harbors a *vanZ* allele, distinct from *vanZ1* (49% identity) present elsewhere in RT106 genome and in other *C. difficile* strains including the well-studied 630 (57). In 630, VanZ1 was previously shown to confer low level resistance to the glycopeptide antibiotic, teicoplanin, but not to vancomycin (57). The presence of a second VanZ allele may contribute to the modest increase in teicoplanin resistance of RT106 strains. The potential selective advantage of this phenotype cannot be ruled out; while teicoplanin is not FDA-approved for use in the US, it is widely used in Europe, Asia and South America.

In addition to the strain 630 cd2831 SrtB-anchored collagen-binding adhesin homolog (99% protein identity) (58), all RT106 strains encode an ortholog within GI1 (locus ID FE556_11350; 33% protein identity with CD2831); this gene was earlier reported as an ‘RT106-associated accessory gene’(29). A subset of RT106 strains (13/94; 6 strains assayed for biofilm formation) also contains an additional ortholog within GI3 (locus ID FE556_02390; 79% protein identity with GI1 locus ID FE556_11350). The robust collagen-dependent biofilm formation observed in the RT106 isolates may be due to the presence of any one or combination of these genes. Further investigation is required to parse the contribution of these genes to virulence. Since toxigenic *C. difficile* can breach the intestinal epithelium via cell junction disruption and/or epithelial cell death, thereby exposing the components of the extracellular matrix including collagen, strong vegetative cell and biofilm adhesion to collagen is a possible mechanism promoting *C. difficile* colonization of, and persistence in, the host.

GI1 also harbors genes for anti-restriction modification (*ardA*), multi-drug resistance (*mfs*), methylglyoxal detoxification (*gloA*), and cobalt-zinc-cadmium resistance protein. Homologs of these genes are linked to virulence of other pathogens (59-61). Finally, GI1 has features of a conjugative mobile element and contains genes for DNA excision/integration and encodes homologs of several proteins involved in Tcp conjugation machinery of *C. perfringens* including TcpC, TcpE, TcpG/TcpI hydrolase, TcpF, and TcpA. It is presently unknown whether the entire 46 kb genomic island can mobilize to other *C. difficile* strains.

Fragments of GI1 were found in different *C. difficile* sequence types. Molecular clock analysis suggests that the complete island is a composite of sequences sequentially acquired from progenitor ST strains. The molecular clock based on the conserved GI1 segment is asynchronous with the one based on house-keeping genes (**Figure 4** and **Supplemental Figure S2**). Further, consistent with higher G/C content of GI1 relative to rest of the *C. difficile* genome, it is likely that the progenitor ST strains acquired the DNA segments from non-clostridial organisms via horizontal gene transfer. While the complete island is yet to be found in any other microbial genome or plasmid, two gene segments (8.4 kb and 13.7 kb) were detected in other enteric bacteria (**Figure 5**). For the 8.4 kb gene segment, the most closely related sequences occur in *Roseburia intestinalis* M50/1 strains (92.2% identity). The 13.7 kb segment displays 99% identity to sequence within a 36 kb genomic island in *E. faecium*. While the 8.4 kb and 13.7 kb segment in GI1 may have been derived from *E. faecium*, the candidate donors of the other gene segments in GI1 are presently unknown. The presence of these genetic segments in disparate enteric organisms may suggest that they confer some selective advantage within the intestinal environment.

## CONCLUSIONS

*C. difficile* RT106 is virulent in a hamster model of infection, and all sequenced isolates within this clade harbor a unique 46 kb GI1. Consistent with the presence of genes encoding a VanZ family protein and a SrtB-anchored collagen-binding adhesin within GI1, RT106 strains had increased teicoplanin resistance and robust collagen-dependent biofilm formation, respectively. Further investigation is required to implicate GI1 genes to RT106 virulence.

## METHODS

### C. difficile surveillance

This study utilized stool samples from clinically-positive, *C. difficile*-infected patients at the Banner University Medical Center (BUMC) in Tucson, Arizona between August 1, 2015 and July 31, 2018. The University of Arizona Institutional Review Board approved this study. All stool samples were de-identified. Samples were collected and stored at −80°C. From August 2015 to February 2017, *tcdB-*positive stool samples tested by BUMC via polymerase chain reaction (PCR) were included in the study. On March 2017, BUMC implemented the glutamate dehydrogenase (GDH) and toxin enzyme immunoassay for *C. difficile* testing. All GDH+ samples were collected. We then screened for the presence of *tcdB* in the GDH+/toxin-samples via PCR using the following primers: B1C (5’-GAAAATTTTATGAGTTTAGTTAATAGAAA-3’) and B2N (5’-CAGATAATGTAGGAAGTAAGTCTATAG-3’) (62). For samples received during March 2017 to July 2018, only GDH+/toxin+ or GDH+/toxin- and *tcdB*-PCR-positive samples were analyzed in this study.

### Ribotyping of clinical C. difficile isolates

Stool samples were plated on taurocholate cycloserine cefoxitin fructose agar (TCCFA) and cultured anaerobically at 37°C. Isolated colonies were lysed with Toothpick™-PCR (G-Biosciences, St. Louis, Missouri, USA), and supernatants were used as templates for ribotyping PCR using the following primers: 16S (5’-GTGCGGCTGGATCACCTCCT-3’) and 23S (5’-CCCTGCACCCTTAATAACTTGACC-3’) (63, 64). Isolated colonies were also submitted to the University of Arizona Genomics Core for genomic extraction using QIAGEN DNeasy column-based extraction kit (QIAGEN, Germantown, Maryland USA) and ribotyping PCR using the same 16S and 23S primers. PCR products were resolved via capillary electrophoresis using an AB Prism® 3730 Genetic Analyzer (Applied Biosystems, Foster City, CA) and amplicon length evaluated using Marker 1.85 (SoftGenetics, State College, PA). Identification of ribotypes from electropherograms was determined using the online resource, Webribo (https://webribo.ages.at/) (63).

### DNA extraction

Genomic DNA samples were extracted using the protocol by Pospiech and Neumann (65), with modifications. Briefly, 50 mL overnight cultures of *C. difficile* were harvested via centrifugation at 3000 g for 10 minutes, and resuspended in 5 mL of SET buffer (75 mM NaCl, 25 mM EDTA, 20 mM Tris, pH 7.5). Cell lysis was facilitated by adding lysozyme (Sigma-Aldrich, St. Louis, MO) to a final concentration of 5 mg/mL and incubating samples at 37°C for 30 minutes. 500 μL of 10% SDS and 25 μL of 100 mg/mL proteinase K (Sigma-Aldrich) were added, and samples were incubated at 55°C for 2 hours. 2.5 mL of 5M NaCl and 5 mL of chloroform (Sigma-Aldrich) was added, and samples mixed with frequent inversions. Samples were centrifuged at 3800 g for 15 minutes, and aqueous phase was collected. DNA was precipitated by 1 volume of isopropanol. DNA was then spooled, transferred to a microfuge tube, rinsed with 70% ethanol, and vacuum dried (Eppendorf, Hauppauge, NY).

### Whole genome sequencing

DNA samples from 38 RT106 samples were submitted to the Office of Knowledge Enterprise Development (OKED) Genomics Core at Arizona State University (Tempe, AZ) for whole genome sequencing. Illumina-compatible genomic DNA libraries were generated on BRAVO NGS liquid handler (Agilent Technologies, Santa Clara, CA) using Kapa HyperPlus KK8514 library kit (Kapa Biosystems, Wilmington, MA). DNA was enzymatically sheared to approximately 600bp fragments, end-repaired and A-tailed as described in the Kapa HyperPlus protocol. Illumina-compatible adapters with unique indexes (IDT #00989130v2; IDT technologies, Skokie, IL) were ligated individually on each sample. The adapter-ligated molecules were cleaned using Kapa pure beads (KK89002, Kapa Biosystems), and amplified with Kapa HiFi DNA Polymerase (KK2502, Kapa Biosystems). Fragment size of each library was analyzed using Agilent Tapestation, and quantified via qPCR using KAPA Library Quantification Kit (KK4835, Kapa Biosystems) and Quantstudio™ 5 Real-time PCR System (Thermo Fisher Scientific, Waltham, MA) before multiplex pooling and sequencing in a 2×250 flow cell on the MiSeq platform (Illumina, San Diego, CA) at the ASU OKED Genomics Core. Genomic libraries were split in 3 MiSeq runs. De novo genome assembly was performed using CLC Genomics Workbench 11 (QIAGEN Bioinformatics, Redwood City, CA). Depth of coverage ranges between 17X-608X (**Supplemental Table S4).** Contigs were annotated via Rapid Annotation using Subsystem Technology (RAST) Version 2.0 (66-68). Sequences for the 38 RT106 genomes were deposited through the National Center for Biotechnology Information (NCBI) Bankit (https://www.ncbi.nlm.nih.gov/WebSub/?tool=genbank) under the GenBank accession numbers listed on Supplemental Table 3.

### Phylogenetic analysis

The 38 RT106 strains were mapped against a collection of all complete or draft *C. difficile* genomes sequences (1425 total sequences) publicly available from the NCBI genome database (January 2019 download date). The phylogenetic tree was generated without sequence alignment by using a Composition Vector approach and CVtree Version 3.0 (30). Interactive Tree of Life v4.3 (https://itol.embl.de/) (69) was used to visualize and annotate the phylogenetic tree.

### In silico multilocus sequence typing (MLST) and in silico ribotyping

Sequence types (ST) of the clinical RT106 isolates and other *C. difficile* strains that claded closely to RT106 in the phylogenetic tree were determined based on the allelic patterns of 7 housekeeping genes (28) using the *C. difficile* MLST database (http://pubmlst.org/cdifficile). In silico ribotyping PCR analysis was performed on the uncharacterized strains using NCBI Primer-Blast (70) and the same 16S and 23S primers listed above. DH/NAP11/106/ST42 (Refseq assembly no. GCF_002234355.1), a complete closed genome, was used as a reference strain for the RT106 PCR fragment pattern.

### Identification of RT106 genomic islands

A series of BLASTN searches (https://blast.ncbi.nlm.nih.gov/Blast.cgi) (71) was performed to identify the unique genetic elements associated with RT106. GV364 sequence was first compared to the complete closed genome sequence of *C. difficile* 630 strain (Refseq assembly no. GCF_000932055.2). Large genetic elements (>10 kb) not found in *C. difficile* 630 were then compared to all 94 RT106 strain sequences to identify genetic elements associated only with RT106. The resulting genetic elements were verified to be unique to RT106 by performing BLASTN searches against 1425 publicly available *C. difficile* genome sequences at the NCBI database.

### Evolutionary Genetic Analysis Using Maximum Likelihood

The 7.1 kb genetic region common to 265 *C. difficile* strains also containing gene segments (>7.7 kb, 98% identity) within the GI1 was used to deduce the possible evolutionary formation of genetic island 1 on RT106. Mega-X (31) was used to perform molecular clock analysis using a maximum likelihood (ML) approach. ML tree was constructed, and branch lengths were adjusted based on Tamura-Nei distance algorithm.

The independence of the acquisition of gene segments forming GI1 was tested by comparing the ML tree to a minimum spanning tree (MST) of MLST allele data profiles in the *C. difficile* MLST database (http://pubmlst.org/cdifficile). MST was created using PhyloViz v2.0 (72).

### Antibiotic susceptibility testing

Overnight cultures of *C. difficile* strains were diluted in brain heart infusion broth (BHI) broth at McFarland scale of 0.5 (approximate OD_600nm_ of 0.1). 100 μL of the culture was plated onto Brucella blood agar. E-test strips (BioMerieux, Durham, NC) for the following antibiotics were applied on the agar: cefotaxime, vancomycin, erythromycin, clindamycin, levofloxacin, metronidazole, moxifloxacin, tetracyline and teicoplanin. Minimum inhibitory concentration, defined as the lowest concentration of the agent that inhibited bacterial growth, was determined. Antibiotic susceptibility was based on Clinical and Laboratory Standard Institute (CLSI) and European Committee on Antimicrobial Susceptibility Testing (EUCAST) breakpoints. There are no set standard breakpoints for teicoplanin.

### Toxin ELISA

Relative levels of TcdA and TcdB toxins were determined using Alere Wampole A/B Toxin ELISA kit (Alere, Atlanta, GA). Overnight cultures of *C. difficile* strains were inoculated in 10 mL BHI at 1:100 dilution. Samples were cultured anaerobically for 72 hours. Cultures were pelleted by centrifugation, and supernatants processed for Toxin ELISA following manufacturer’s protocol and using BioTek Synergy automated plate reader (Bio-Tek, Winooski, VT). Total protein present in the supernatants were quantified using Pierce BCA protein assay kit (Thermo Fisher Scientific, Waltham, MA). Relative amounts of toxin were normalized to total proteins.

### Motility assay

Motility agar plates were prepared by adding 20 mL of BHI with 0.3% agar per well of a Corning Costar 6-well plate (Thermo Fisher Scientific, Waltham, MA). *C. difficile* strains were cultured in BHI overnight. Approximately 5 μL of the culture was collected and stabbed into the motility agar. Plates were sealed and incubated in a humid, anaerobic chamber for 72 hours, and then imaged using ChemiDoc™ Touch Imaging System (Bio-rad, Hercules, CA).

### Biofilm assay

Twenty-four well plates were coated with: human or rat tail collagen type I (88 ng per well; Sigma-Aldrich), human collagen type III (88 ng per well; Sigma-Aldrich) or a combination of human collagen type I and type III (88 ng of each collagen type per well). Overnight cultures of *C. difficile* strains were diluted in BHIS containing 100 mM glucose (OD_600nm_=0.1). One mL of the culture was added per well of the uncoated or collagen-coated plate and incubated anaerobically for 72 hours at 37°C. Supernatants were removed gently by tilting plates onto a collection basin. Biofilms were washed twice by gently submerging plates in glass basins of PBS. Excess PBS was removed by inverting plates onto tissue paper. Biofilms were fixed for 20-40 minutes at 37°C, and then stained with 1 mL of 0.2% filter-sterilized crystal violet for 30 minutes. Biofilms were washed twice with PBS as described above. For quantification of biofilm growth, 1mL of 4:1 ethanol/acetone solution was added to each sample. 100 μL aliquots were transferred to a 96-well plate, and absorbance at 570 nm (A_570nm_) was determined using BioTek Synergy automated plate reader (BioTek Instruments, Inc., Winooski, VT). Relative changes in biofilm densities (ΔA_570nm_) were determined by comparing A_570nm_ of crystal violet-stained biofilms formed on collagen-coated vs. on uncoated plastic wells.

### C. difficile infection of Golden Syrian hamsters

A pilot hamster study, approved by the Institutional Animal Care and Use Committee of the University of Arizona, was conducted to test the virulence of GV599. Six-week old male Golden Syrian hamsters (weighing 90-110 grams) procured from Charles River Laboratory (Wilmington, MA) were given clindamycin and chloramphenicol prior to infection with *C. difficile* spores. Clindamycin (prescription solution; clindamycin sulfate; University of Arizona Pharmacy; 30 mg/kg) was administered as a single oral dose three days prior to infection (Day -3). Chloramphenicol (Sigma-Aldrich; 50 mg/kg) was administered orally for a total of 6 doses prior to infection (2 doses on Day -3, 3 doses on Day -2, and 1 dose on Day -1). Three hamsters were infected with GV599 (164 spores; orally administered in PBS). The control hamster was given both antibiotics and PBS. Animals were monitored for disease symptoms (wet tail, ruffled coat, lethargy, weight loss) through the course of the study. Moribund hamsters or those meeting the criteria for euthanasia were administered 270 mg/kg Euthanasia III (MedPharma Inc, Pomona, CA, United States). Euthanized hamsters were dissected for visualization of gross pathology, and cecal contents harvested and plated on selective TCCFA to confirm *C. difficile* colonization. Colonies recovered were ribotyped for confirmation of RT106 infection. Cecal tissue samples were fixed with 10% neutral buffered formalin and submitted to Arizona Veterinary Diagnostic Laboratory (Tucson, AZ) for hematoxylin and eosin staining.

## LIST OF ABBREVIATIONS

CDI: (*Clostridioides difficile* infections),
CDC: (Center for Disease Control and Prevention),
BUMC: (Banner University Medical Center),
MLST: (multi-locus sequence typing),
CLSI: (Clinical and Laboratory Standard Institutes),
PaLoc: (pathogenicity locus),
RT: (ribotype),

## DECLARATIONS

1. Ethics approval and consent to participate
2. Consent for publication
3. Availability of data and material The datasets generated and/or analyzed during the current study are available in the GenBank repository (BioProject PRJNA542726).
4. Competing interests
  ○ The authors declare that they have no competing interests
5. Funding This work was supported by funding from the National Institutes of Health [R33AI121590531(GV) and the US Dept. of Veterans Affairs [IK6BX003789(GV); I01BX001183(GV)].
6. Authors’ contributions BPR performed the comparative genomic analyses. JLR performed phenotypic characterization and statistical analysis, analyzed data and drafted the manuscript. RCW, ASM and FA isolated *C. difficile* from stool samples and performed ribotyping analysis. AH performed moxifloxacin and teicoplanin MIC determination and biofilm assay experiments. AW, JL and SJ conducted the pilot hamster experiment. SPE and KWS contributed in the conceptualization of the research and analyzed data. GV and VKV conceptualized and funded the studies, finalized the manuscript and provided full project oversight. All authors read and approved the manuscript.
7. Authors’ information (optional)

