## Supplemental Data for "Phylogenomic analysis of *Clostridiodes difficile* ribotype 106 strains reveals novel genetic islands and emergent phenotypes"

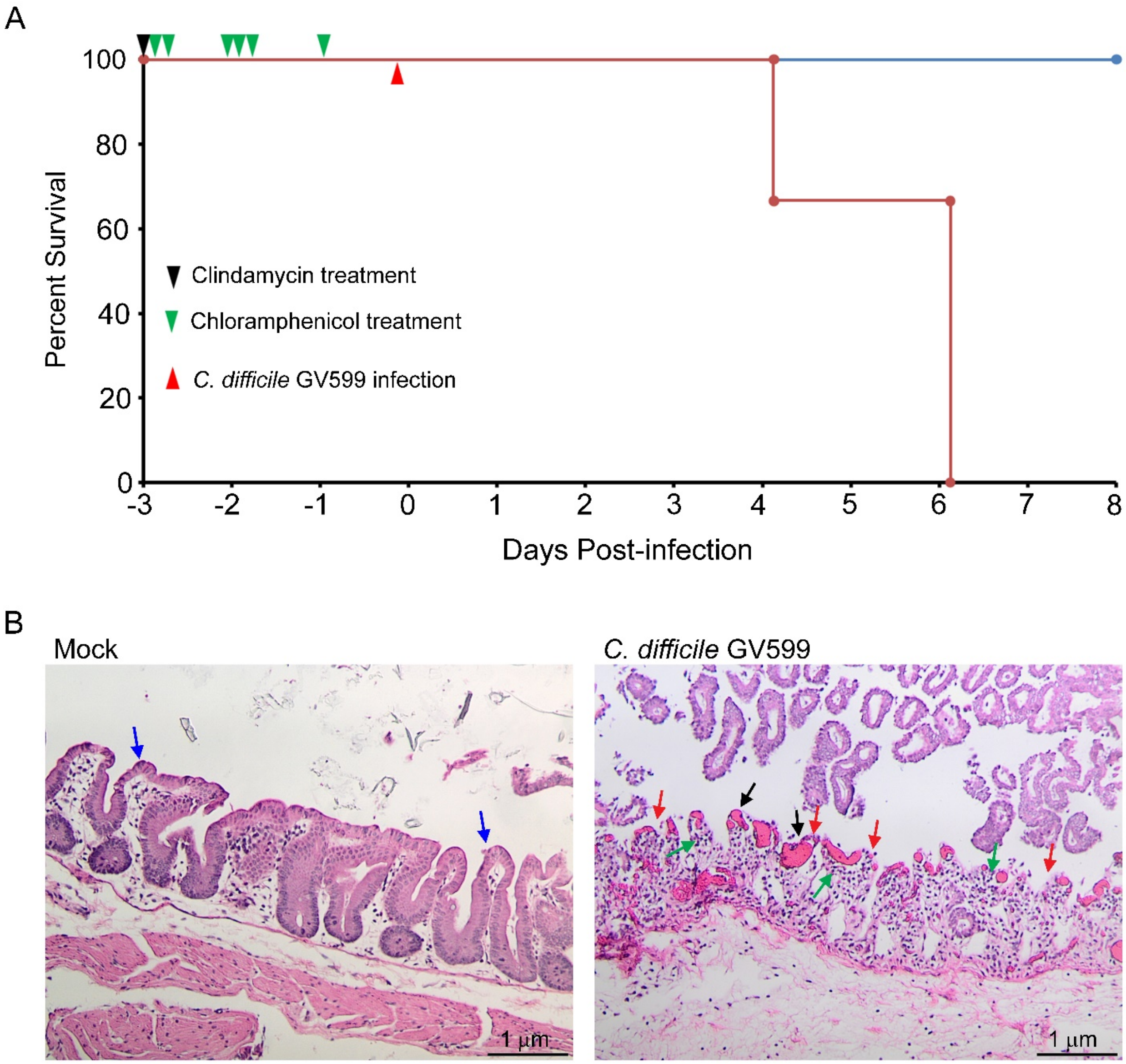


**Supplemental Figure S1. Golden Syrian hamsters succumbed to *C. difficile* GV599 infection. A**, Infection of Golden Syrian hamsters with GV599 (164 spores in PBS; administered orally) caused 100% mortality (red line; n=3). Animals were treated with a single dose of clindamycin (30mg/kg body weight) three days prior to infection (blue arrowhead) and a total of six doses of chloramphenicol (50mg/kg body weight)

administered on Days -3, -2 and -1 prior to infection (green arrowhead). The study was ended two days after the last GV599-infected hamster died, at which point the mock- or PBS-treated hamster (n=1) was euthanized. **B**, Microscopic images of hematoxylin-eosin-stained colonic tissues of GV599-infected hamsters revealed classic *C. difficile* infection pathology including gross hemorrhage (black arrows), epithelial erosion (red arrow) and recruitment of inflammatory infiltrates (green arrows). In comparison, colonic tissue sections of the mock-treated hamster had intact intestinal epithelium (blue arrow). Images were captured using an Olympus SC30 microscope equipped with a DPlan 10X 0.25 160/0.17 objective lens (Olympus, Center Valley, PA), and are representative of 27 fields of view per animal.

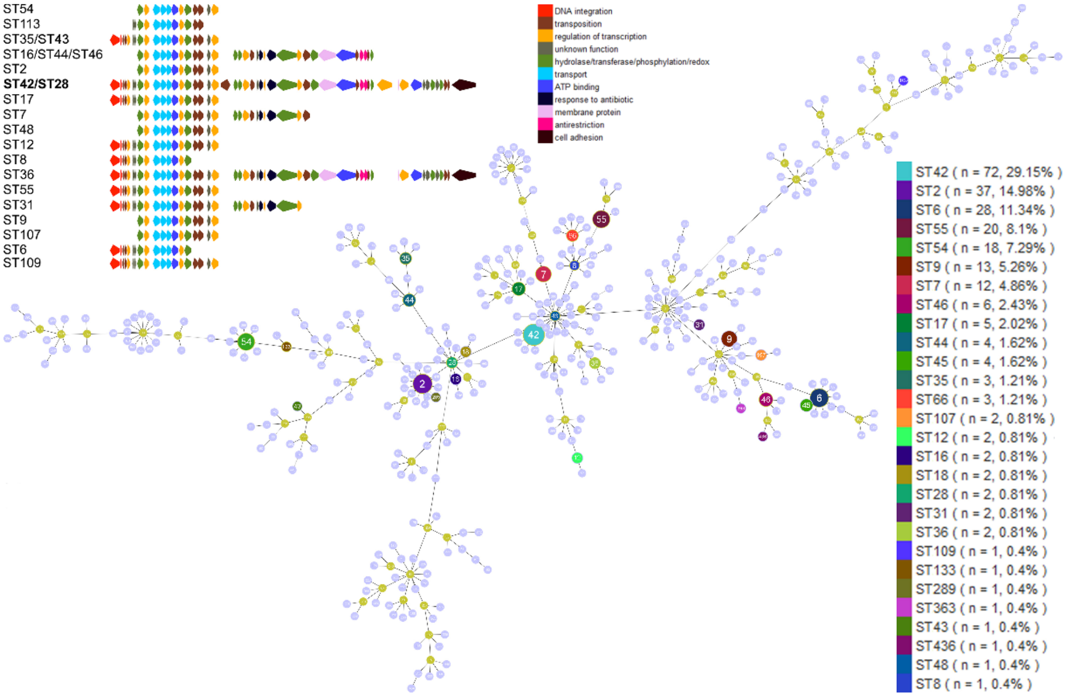


**Supplemental Figure S2. Sequence type does not recapitulate carriage of the 46 kb island 1.** Minimum spanning tree (MST) of the *C. difficile* sequence types (ST) that carry the whole or part of the 46 kb genomic island 1. Analysis was performed on MLST allele data from PubMLST (https://pubmlst.org/). Tree shows relatedness of the STs independent of genomic island 1.

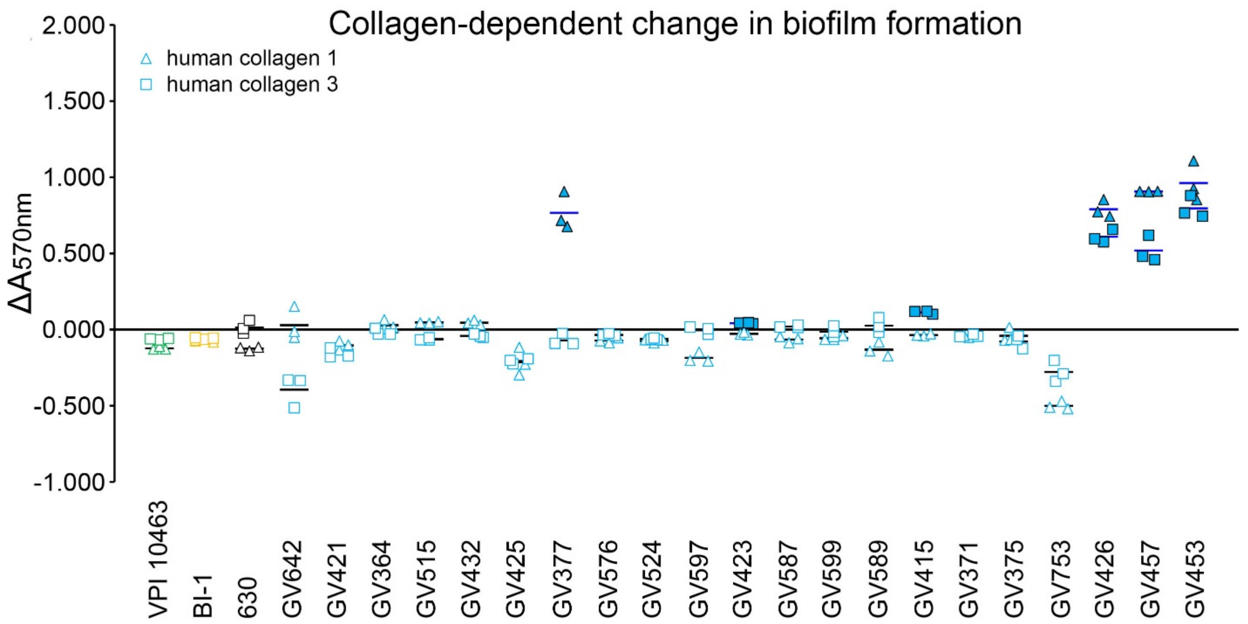


**Supplemental Figure S3. Clinical RT106 isolates form dense biofilms on human type I and type III collagen.** 21 clinical RT106 strains (blue) and 3 non-RT106 toxigenic *C. difficile* strains (VPI, BI-1, and 630 designated as green, yellow and black, respectively) were cultured for 72 hours in uncoated wells or wells coated with human type I or type III collagen. Biofilm assay was performed by measuring absorbance at $\lambda$=570 nm ($A_{570nm}$) of solubilized crystal violet-stained biofilms. Relative changes in biofilm densities ($\Delta A_{570nm}$) were determined by comparing $A_{570nm}$ of crystal violet-stained biofilms formed on human type I or type III collagen vs. on uncoated plastic wells. Triangles denote $\Delta A_{570nm}$ for human type I collagen, while squares denote $\Delta A_{570nm}$ for human type III collagen. Student's t test was performed to compare mean absorbance at 570nm of biofilm formed by each strain on collagen-coated vs. uncoated wells. Filled blue triangles or squares denote $P_{value} < 0.05$. No difference in biofilm formation was observed when the reference *C. difficile* 630, BI-1 and VPI strains were cultured on wells with or without collagen. One-sample one-tailed T-test was performed to determine whether the group of RT106 strains displayed denser biofilms on collagen-coated wells ($H_{alt}$: mean $\Delta A_{570nm}>0$; $H_0$: mean $\Delta A_{570nm}=0$; $P_{value}=0.13903$ for type I collagen data set; $P_{value}=0.305584$ for type III collagen data set).

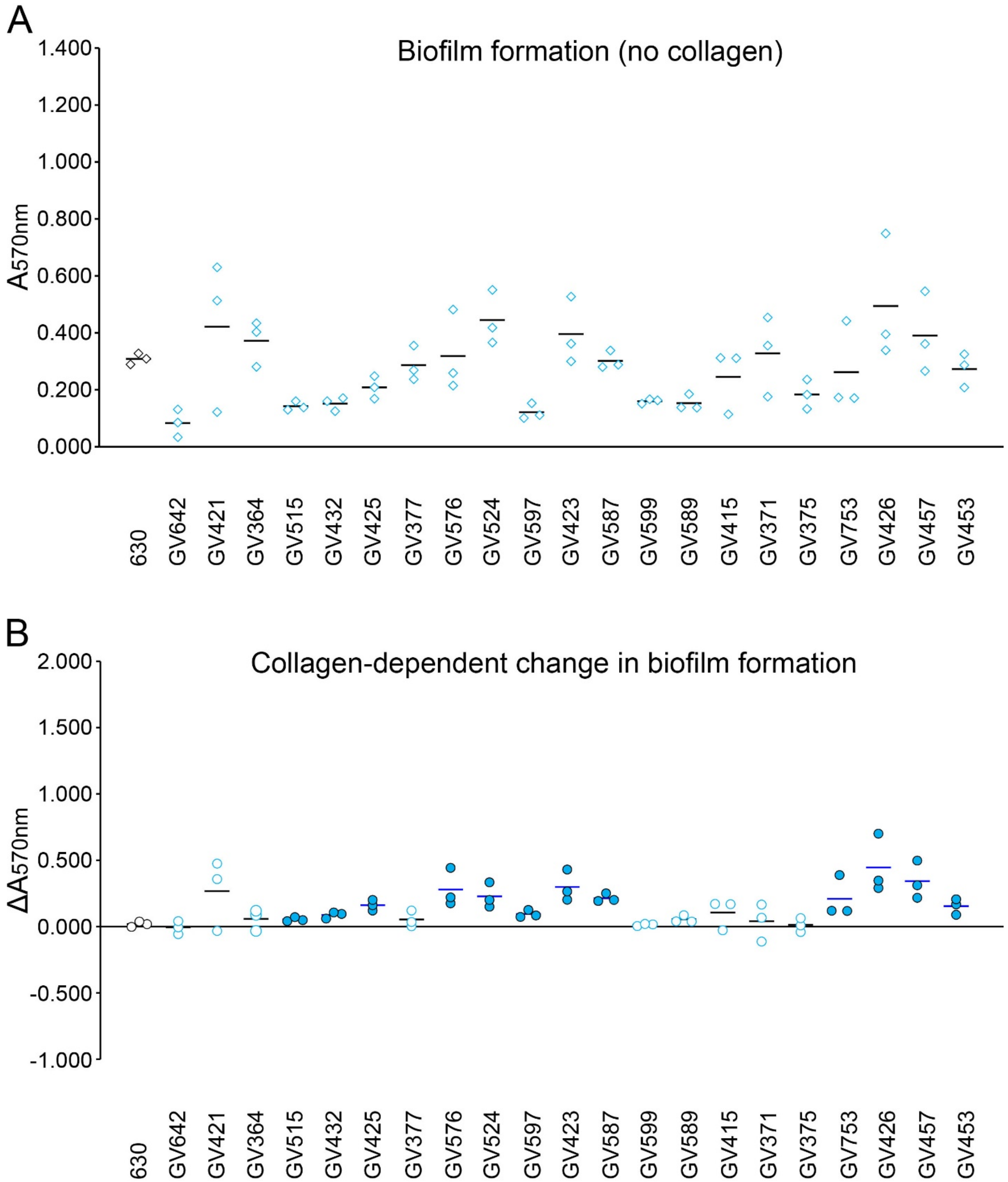


**Supplemental Figure S4. Clinical RT106 isolates form dense biofilms on rat tail type I collagen. A**, 21 clinical RT106 strains (blue) and a non-RT106 toxigenic *C. difficile* 630 strain (black) were cultured for 72 hours in uncoated wells or wells coated with human type I or type III collagen. Biofilm assay was performed by measuring absorbance at $\lambda$=570 nm ($A_{570nm}$) of solubilized crystal violet-stained biofilms. RT106 isolates displayed variable biofilm densities on abiotic plastic wells. **B,** Relative changes in biofilm densities ($\Delta A_{570nm}$) were determined by comparing $A_{570nm}$ of crystal violet-stained biofilms formed on rat tail type I collagen vs. on uncoated plastic wells. Student's t test was performed to compare mean absorbance at 570nm of biofilm formed by each strain on collagen-coated vs. uncoated wells. Filled blue circles denote $P_{value} < 0.05$. No difference in biofilm formation was observed when the reference *C. difficile* 630 was cultured on wells with or without collagen. One-sample one-tailed T-test was performed to determine whether the group of RT106 strains displayed denser biofilms on collagen-coated wells ($H_{alt}$: mean $\Delta A_{570nm}>0$; $H_0$: mean $\Delta A_{570nm}=0$; $P_{value}=0.00001$).

### Supplemental Table S1. List of genes within genomic island 1

| Locus ID | Product | Start | End | Strand |
|---|---|---|---|---|
| FE556_11090 | site-specific integrase | 201569 | 202759 | - |
| FE556_11095 | excisionase | 202837 | 203040 | - |
| FE556_11100 | hypothetical protein | 203145 | 203382 | + |
| FE556_11105 | helix-turn-helix domain-containing protein | 203412 | 203666 | - |
| FE556_11110 | sigma-70 family RNA polymerase sigma factor | 203669 | 204088 | - |
| FE556_11115 | hypothetical protein | 204355 | 204585 | - |
| FE556_11120 | DUF4177 domain-containing protein | 204609 | 204800 | - |
| FE556_11125 | class I SAM-dependent methyltransferase | 204964 | 205680 | - |
| FE556_11130 | TetR/AcrR family transcriptional regulator | 205802 | 206428 | - |
| FE556_11135 | cation transporter | 206910 | 207821 | - |
| FE556_11140 | ABC transporter permease | 207833 | 208531 | - |
| FE556_11145 | ABC transporter permease | 208528 | 209211 | - |
| FE556_11150 | ABC transporter ATP-binding protein | 209208 | 210053 | - |
| FE556_11155 | TetR/AcrR family transcriptional regulator | 210157 | 210744 | - |
| FE556_11160 | alpha/beta hydrolase | 210816 | 211673 | - |
| FE556_11165 | conjugal transfer protein | 211886 | 212497 | - |
| FE556_11170 | bifunctional lytic transglycosylase/NlpC/P60 family protein | 212448 | 213188 | - |
| FE556_11175 | DUF3788 domain-containing protein | 213656 | 214090 | - |
| FE556_11180 | helix-turn-helix domain-containing protein | 214168 | 215079 | - |
| FE556_11185 | IS110 family transposase | 215358 | 216527 | + |
| FE556_11190 | GNAT family N-acetyltransferase | 216917 | 217468 | - |
| FE556_11195 | SAM-dependent methyltransferase | 217482 | 217958 | - |
| FE556_11200 | MerR family transcriptional regulator | 218039 | 218854 | - |
| FE556_11205 | conjugal transfer protein | 218957 | 219529 | - |
| FE556_11210 | DUF4865 family protein | 219799 | 220074 | - |
| FE556_11215 | VanZ family protein | 220079 | 220585 | - |
| FE556_11220 | XRE family transcriptional regulator | 220688 | 220881 | + |
| FE556_11225 | tetracycline resistance MFS efflux pump | 221068 | 222231 | - |
| FE556_11230 | pyruvate, phosphate dikinase | 222221 | 224788 | - |
| FE556_11235 | TetR/AcrR family transcriptional regulator | 224785 | 225396 | - |
| FE556_11240 | conjugal transfer protein | 225540 | 226442 | - |
| FE556_11245 | bifunctional lytic transglycosylase/NlpC/P60 family protein | 226462 | 227466 | - |
| FE556_11250 | MFS transporter | 227463 | 229658 | - |
| FE556_11255 | ATP-binding protein | 229658 | 232108 | - |
| FE556_11260 | conjugal transfer protein | 232086 | 232484 | - |
| FE556_11265 | antirestriction protein ArdA | 232605 | 233108 | - |
| FE556_11270 | antirestriction protein ArdA | 233126 | 233629 | - |
| FE556_11275 | conjugal transfer protein | 233548 | 233844 | - |
| FE556_11280 | GNAT family N-acetyltransferase | 233951 | 234331 | - |
| FE556_11285 | hypothetical protein | 234409 | 234570 | - |
| FE556_11290 | group II intron reverse transcriptase/maturase ltrA | 234737 | 236647 | - |
| FE556_11295 | DUF3789 domain-containing protein | 237389 | 237523 | - |
| FE556_11300 | replication initiation factor domain-containing protein | 237530 | 238651 | - |
| FE556_11305 | ATP-binding protein | 238931 | 240325 | - |
| FE556_11310 | YcxB family protein | 240436 | 240747 | - |
| FE556_11315 | DUF3795 domain-containing protein | 240775 | 241317 | - |
| FE556_11320 | alpha/beta hydrolase | 241378 | 241980 | - |
| FE556_11325 | DUF3788 domain-containing protein | 242035 | 242448 | - |
| FE556_11330 | glyoxalase | 242512 | 242970 | - |
| FE556_11335 | DUF961 domain-containing protein | 243051 | 243470 | - |
| FE556_11340 | DUF961 domain-containing protein | 243479 | 243802 | - |
| FE556_11345 | hypothetical protein | 243759 | 244011 | - |
| FE556_11350 | SrtB-anchored collagen-binding adhesin | 244024 | 247071 | - |

**Supplemental Table S2. List of genes within genomic island 3**

| Locus ID | Product | Start | Stop | Stand |
|---|---|---|---|---|
| FE556_02390 | SrtB-anchored collagen-binding adhesin | 145981 | 149034 | + |
| FE556_02395 | DNA cytosine methyltransferase | 149035 | 150111 | + |
| FE556_02400 | DUF961 domain-containing protein | 150312 | 150635 | + |
| FE556_02405 | DUF961 domain-containing protein | 150652 | 151020 | + |
| FE556_02410 | ATP-binding protein | 151038 | 152435 | + |
| FE556_02415 | XRE family transcriptional regulator | 152703 | 153929 | + |
| FE556_02420 | DUF3789 domain-containing protein | 153913 | 154056 | + |
| FE556_02425 | hypothetical protein | 154057 | 154278 | + |
| FE556_02430 | iron-sulfur protein | 154352 | 155113 | + |
| FE556_02435 | conjugal transfer protein | 155212 | 155433 | + |
| FE556_02440 | antirestriction protein ArdA | 155430 | 155915 | + |
| FE556_02445 | antirestriction protein ArdA | 155932 | 156435 | + |
| FE556_02450 | conjugal transfer protein | 156521 | 156910 | + |
| FE556_02455 | ATP-binding protein | 156897 | 159347 | + |
| FE556_02460 | YtxH domain-containing protein | 159410 | 161512 | + |
| FE556_02465 | peptidase P60 | 161509 | 162519 | + |
| FE556_02470 | conjugal transfer protein | 162537 | 163448 | + |
| FE556_02475 | TetR/AcrR family transcriptional regulator | 163589 | 164215 | + |
| FE556_02480 | ABC transporter ATP-binding protein | 164212 | 165099 | + |
| FE556_02485 | ABC transporter permease | 165096 | 165893 | + |
| FE556_02490 | ABC transporter ATP-binding protein | 166073 | 166759 | + |
| FE556_02495 | ABC transporter permease | 166749 | 169307 | + |
| FE556_02500 | response regulator transcription factor | 169372 | 170037 | + |
| FE556_02505 | HAMP domain-containing histidine kinase | 170040 | 171065 | + |
| FE556_02510 | sigma-70 family RNA polymerase sigma factor | 171349 | 171771 | + |
| FE556_02515 | helix-turn-helix domain-containing protein | 171776 | 172015 | + |
| FE556_02520 | phosphoesterase | 172468 | 173319 | + |
| FE556_02525 | DUF4368 domain-containing protein | 173612 | 175225 | + |

**Supplemental Table S3. Increase in minimum inhibitory concentration after sub-inhibitory exposure to teicoplanin**

| Strain | Fold change in teicoplanin resistance post-induction |
|---|---|
| GV415 | + 1.33 |
| GV425 | + 2.00 |
| GV432 | + 1.33 |
| GV457 | + 1.52 |
| GV576 | + 1.47 |
| GV589 | + 1.47 |
| GV753 | + 4.04 |

### Supplemental Table S4. GenBank Accession numbers of RT106 isolates

| Strain | NCBI Accession Numbers | | | Coverage |
|---|---|---|---|---|
| | Biosample | BioProject | WGS | |
| GV364 | SAMN11637479 | PRJNA542726 | VCAN00000000 | 608x |
| GV371 | SAMN11637480 | PRJNA542726 | VCAM00000000 | 463x |
| GV375 | SAMN11637481 | PRJNA542726 | VCAL00000000 | 543x |
| GV377 | SAMN11637482 | PRJNA542726 | VCAK00000000 | 550x |
| GV415 | SAMN11637483 | PRJNA542726 | VCAJ00000000 | 383x |
| GV421 | SAMN11637484 | PRJNA542726 | VCAI00000000 | 476x |
| GV423 | SAMN11637485 | PRJNA542726 | VCAH00000000 | 414x |
| GV425 | SAMN11637486 | PRJNA542726 | VCAG00000000 | 436x |
| GV426 | SAMN11637487 | PRJNA542726 | VCAF00000000 | 594x |
| GV432 | SAMN11637488 | PRJNA542726 | VCAE00000000 | 512x |
| GV453 | SAMN11637490 | PRJNA542726 | VCAC00000000 | 163x |
| GV457 | SAMN11637489 | PRJNA542726 | VCAD00000000 | 146x |
| GV515 | SAMN11637491 | PRJNA542726 | VCAB00000000 | 161x |
| GV524 | SAMN11637492 | PRJNA542726 | VCAA00000000 | 89x |
| GV576 | SAMN11637493 | PRJNA542726 | VBZZ00000000 | 101x |
| GV587 | SAMN11637496 | PRJNA542726 | VBZW00000000 | 110x |
| GV589 | SAMN11637497 | PRJNA542726 | VBZV00000000 | 78x |
| GV597 | SAMN11637495 | PRJNA542726 | VBZX00000000 | 117x |
| GV599 | SAMN11637494 | PRJNA542726 | VBZY00000000 | 114x |
| GV642 | SAMN11637498 | PRJNA542726 | VBZU00000000 | 91x |
| GV753 | SAMN11637499 | PRJNA542726 | VBZT00000000 | 126x |
| GV814 | SAMN11637501 | PRJNA542726 | VBZR00000000 | 30x |
| GV831 | SAMN11637500 | PRJNA542726 | VBZS00000000 | 28x |
| GV836 | SAMN11637502 | PRJNA542726 | VBZQ00000000 | 32x |
| GV840 | SAMN11637503 | PRJNA542726 | VBZP00000000 | 32x |
| GV868 | SAMN11637504 | PRJNA542726 | VBZO00000000 | 33x |
| GV870 | SAMN11637505 | PRJNA542726 | VCDT00000000 | 25x |
| GV962 | SAMN11637506 | PRJNA542726 | VBZN00000000 | 34x |
| GV973 | SAMN11637507 | PRJNA542726 | VBZM00000000 | 33x |
| GV986 | SAMN11637508 | PRJNA542726 | VBZL00000000 | 31x |
| GV996 | SAMN11637509 | PRJNA542726 | VBZK00000000 | 34x |
| GV997 | SAMN11637510 | PRJNA542726 | VBZJ00000000 | 22x |
| GV1002 | SAMN11637511 | PRJNA542726 | VBZI00000000 | 27x |
| GV1006 | SAMN11637512 | PRJNA542726 | VBZH00000000 | 30x |
| GV1057 | SAMN11637513 | PRJNA542726 | VBZG00000000 | 36x |
| GV1105 | SAMN11637514 | PRJNA542726 | VBZF00000000 | 17x |
| GV1125 | SAMN11637515 | PRJNA542726 | VBZE00000000 | 24x |
| GV1152 | SAMN11637516 | PRJNA542726 | VBZD00000000 | 35x |
